# Abrupt and gradual changes in neuronal processing upon falling asleep and awakening

**DOI:** 10.1101/2024.02.06.579189

**Authors:** Amit Marmelshtein, Barak Lavy, Barak Hadad, Yuval Nir

**Author notes:** These authors contributed equally (Amit Marmelshtein, Barak Lavy). Corresponding author: Yuval Nir.

## Abstract

The neural processes that change when falling asleep are only partially understood. At the cortical level, features of both spontaneous neural activity and sensory responses change between wakefulness and sleep. For example, in early auditory cortex, sleep increases the occurrence of post-onset silent (OFF) periods and elevates population synchrony. However, it remains unknown whether such changes occur abruptly or gradually around sleep onset and awakening. Here, we recorded spontaneous and sound-evoked neuronal spiking activity in early auditory cortex along with polysomnography during thousands of episodes when male rats fell asleep or woke up. We found that when falling asleep, stimulus-induced neuronal silent periods (OFF periods), characteristic of non-rapid eye movement (NREM) sleep, increased within few seconds around sleep onset. By contrast, a gradual increase in neuronal population synchrony built up over tens of seconds until reaching maximal levels. EEG auditory-evoked potentials likely representing stimulus-triggered “K complexes” changed along with post-onset neuronal firing, whereas ongoing EEG slow wave activity was associated with neuronal population synchrony. Similar effects, but with opposite direction, were observed around awakenings. The results highlight late stimulus-induced neuronal silence as a key feature changing abruptly around transitions between vigilance states, likely reflecting neuronal bistability and manifesting also in EEG evoked potentials. More generally, these findings emphasize the added value of going beyond monitoring ongoing activity and perturbing the nervous system to reveal its state - an insight that could also help guide development of more sensitive non-invasive monitors of falling asleep in humans.

## Introduction

Every night we drift into sleep, becoming disconnected as our conscious experience of the sensory environment largely subsides, raising fascinating questions regarding the ways in which brain states shape our behavior and subjective experiences. Previous work has investigated how different brain states affect sensory processing at the neuronal level (Harris and Thiele, 2011; Lee and Dan, 2012; Mcginley et al., 2015). In the auditory domain, such research revealed how factors such as arousal, locomotion, sleep pressure, ongoing behavioral tasks, and attention influence auditory responses (Edeline et al., 2001; Issa and Wang, 2008; Atiani et al., 2009; Harris and Thiele, 2011; Nir et al., 2013; Sela et al., 2020; Marmelshtein et al., 2023).

In a recent investigation, we examined responses to sounds within the early auditory cortex of rats across vigilant states. Our findings revealed robust modulation in specific aspects of neuronal activity, particularly in post-onset responses, entrainment to fast click-trains, and population synchrony (Marmelshtein et al., 2023). Population synchrony, in this context, was defined as “population coupling” (Okun et al., 2015), a measure of how correlated each unit’s firing is with the firing of the local population, and post-onset response as the firing rate within the [30,80]ms interval following stimulus onset (and onset response). In contrast, spontaneous firing rates only showed modest modulation, while onset responses were largely unaffected (Marmelshtein et al., 2023). While these changes were examined over dozens of minutes or even hours, abrupt brain state changes accompanying sleep onset and offset offer a valuable model for understanding how changes in brain states can rapidly influence behavior and perception beyond factors operating at slower time scales.

Falling asleep reflects a progressing sequence of changes in electrical, physiological, behavioral, and phenomenological phenomena (Ogilvie, 2001; Lacaux et al., 2023) governed by changes in subcortical wake- and sleep-promoting neuromodulation (Nir and de Lecea, 2023). Changes in heart rate, blood flow, eye movements, behavioral reaction times, as well as reports of subjective experience take minutes to develop and coincide with sequential changes in electroencephalogram (EEG) activity. Such changes comprise diminished alpha and increased cortical theta activity, occurrence of sharp-waves, and ultimately include bona-fide sleep events such as K-Complexes and sleep spindles (Hori et al., 1994; Adamantidis et al., 2019). These changes represent an inherently gradual and continuous process, not occurring in any single moment (Ogilvie, 2001). On the other hand, measures of behavioral responsiveness, and possibly some EEG phenomena, do change rapidly within few seconds around sleep onset (Ogilvie, 2001; Strauss et al., 2022). Is there then a functionally-relevant abrupt binary switch within the extended process of falling asleep? We sought to address this open question by testing whether changes in cortical neuronal activities accompanying sleep onset exhibit abrupt or gradual dynamics.

To date, sleep onset studies with high temporal resolution (seconds) were typically performed non-invasively in humans, focusing on timing and topographical changes in different EEG components and functional magnetic resonance imaging (fMRI) measurements (Hori et al., 1994; Fukunaga et al., 2006). Auditory EEG responses around sleep onset reveal that signatures of low-level auditory processing, as reflected in early EEG components or responses related to stimulus-specific adaptation, are not robustly modulated by sleep onset (Strauss et al., 2015). By contrast, measures of high-level processing such as P300, responses linked to predictive coding, and measures of baseline EEG complexity and connectivity, are strongly suppressed upon falling asleep (Strauss et al., 2015, 2022). However, non-invasive studies lack the ability to investigate processes at the neuronal level. On the other hand, studies focusing on cortical sensory responses at the neuronal level (Issa and Wang, 2008, 2011, 2013; Sela et al., 2016, 2020; Hayat et al., 2022; Marmelshtein et al., 2023) have not investigated modulations within seconds around state transitions. Here, we aimed to bridge this gap by investigating, with high-temporal resolution, changes in neuronal activity in the auditory cortex as rats fell asleep or woke up. Our findings reveal that post-onset silence (or firing) undergoes substantial abrupt changes within seconds upon falling asleep (or awakening). Other aspects of auditory processing such as elevated population synchrony, vary at a much slower time scale, reflecting gradual aspects of falling asleep. We also find rapid modulation in frontal EEG activity exhibiting a selective association between the EEG sound-evoked potential (likely reflecting a stimulus-induced K-complex) and the auditory cortex post-onset response, whereas baseline slow wave activity is associated with auditory cortex population synchrony.

## Results

We investigated how sensory processing changes during the transitions between wakefulness and sleep by conducting sixteen experiments on adult male Wister rats (n = 6) implanted with microwire arrays targeting the auditory cortex, as well as monitoring with polysomnography including electroencephalography (EEG), electromyography (EMG), and synchronized video (Methods, Figure 1A-B). Rats were situated within a computer-controlled motorized running wheel placed inside an acoustic chamber for a total duration of 10 hours, commencing at light onset. In the first part of the experiment (5 hours), the wheel was periodically rotated for 4-7 seconds each time, alternating with idle intervals of 36-39 seconds (Figure 1D). In the second part of the experiment (recovery), rats were allowed to sleep undisturbed (while still in the wheel apparatus) for an additional 5 hours, during which they naturally transitioned between sleep and wakefulness. Throughout the entire 10h experimental period, we continuously presented auditory stimuli, 500ms-long click-trains at 2, 10, 20, 30 and 40Hz (Figure 1C) with an average inter-stimulus interval of 2s, played from an overhead free-field speaker (Methods) so that many of these sounds occurred around episodes of falling asleep or awakenings.

**Figure 1.**
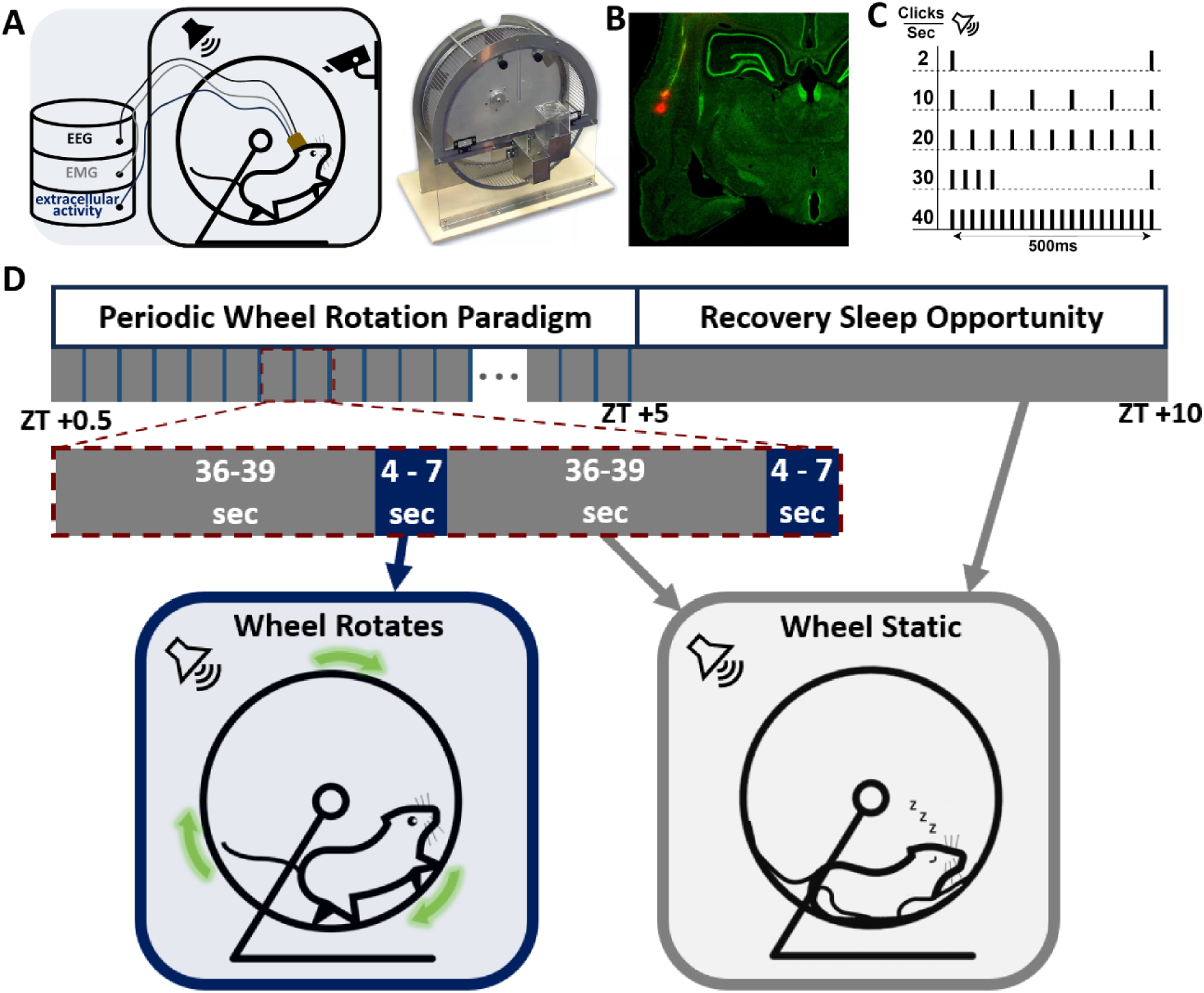
Experimental Setup. (A) (left): Acoustic chamber with an ultrasonic speaker and video monitoring, in which rats were confined to a motorized running wheel, while EEG, EMG, and extracellular neural activity were measured continuously. (right): Motorized running wheel (Lafayette instrument), to which the rats were confined. (B) Histology of microwire traces targeting the auditory cortex. (C) Click-trains at varying rates presented throughout the experiment (D) Schematic depiction of the experimental paradigm. For the first 5h (zeitgeber time [ZT] 0–5), rats were intermittently forced to perform a short running bout with breaks allowing them to fall asleep. It was followed by 5h of undisturbed recovery sleep opportunity (zeitgeber time [ZT] 5–10). Auditory stimuli were presented throughout the entire experiment.

In the first 5h part of the experiment (periodic wheel rotation paradigm), rats often (44.3%) fell asleep during idle intervals and awakened with each wheel rotation, thereby presenting numerous isolated episodes of falling asleep (n=2,971 total, 186 ± 12.1 per session in n=16 sessions, mean ± SEM, Figure 2A,B,D). The second 5h part of the experiment (recovery sleep opportunity) also resulted in a substantial number of falling asleep episodes (n=2,996 total, 187± 16.6 per session, mean ± SEM, Figure 2A,C,D), as well as many spontaneous awakenings (n=2,718 total, 170± 17.2 per session, mean ± SEM, Figure 2A,D). In total, data analysis was performed for 5967 episodes of falling asleep and 2718 episodes of awakening.

**Figure 2.**
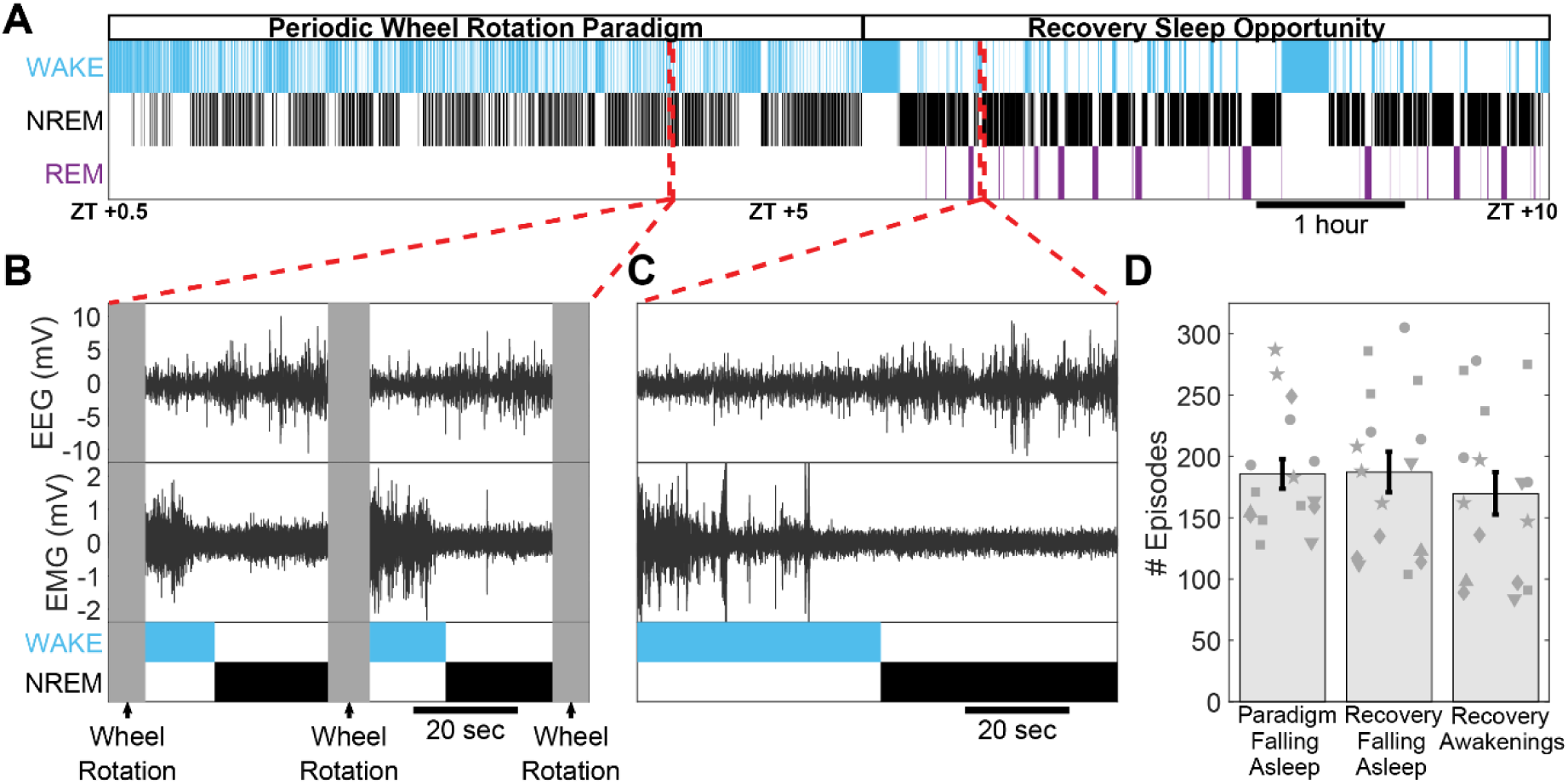
Capturing moments of falling asleep with polysomnography. (A) Representative hypnogram of a typical 10h session. Rows mark epochs of wakefulness (cyan), NREM sleep (black), and REM sleep (purple). Red dashed lines mark two example moments of a rat falling asleep – either between wheel rotations or during subsequent recovery sleep. (B) Representative Polysomnography of a trial of a rat falling asleep during two consecutive 40s intervals between brief (5s) wheel rotations (vertical gray bars). Rows (top to bottom) mark EMG, EEG, and automatically scored vigilance state (top – wake, cyan; bottom – NREM sleep, black) as in panel A. (C) Same as B, but for an example of a rat falling asleep during the recovery sleep opportunity period. (D) Count distribution across all sessions (n = 16) of falling-asleep instances during the periodic wheel rotation paradigm, as well as falling-asleep instances and awakenings during the recovery sleep opportunity period (left to right, respectively).

### Identifying episodes of falling asleep and awakening

We first set out to automatically identify each falling asleep episode. To this end, we implemented the neural network-based automated AccuSleep algorithm (Barger et al., 2019) (Methods, Figure 3). The algorithm was trained on 1h-long manually-scored recovery period data containing intervals of wakefulness, non-rapid eye movement (NREM) sleep, and rapid eye movement (REM) sleep. We focused on the first hour of the recovery sleep opportunity that likely represents the deepest NREM sleep, in an effort to take a conservative approach in recognizing NREM and falling asleep episodes that refrains from subsequent analysis of borderline cases. To evaluate the performance of the automatic sleep-scoring method, we analyzed three sessions with full manual sleep scoring (∼10 hours each). We found that 93 ± 1.1% of automatically scored 1-second epochs correctly matched their corresponding manually scored labels (Figure 3D). This resulted in closely aligned sleep onset timings, with mean ± standard deviation time differences of -1.5 ± 4.6 s, 0.1 ± 5 s, and 2.8 ± 4.4 s between automatic and manual scoring across the three sessions (Methods).

**Figure 3.**
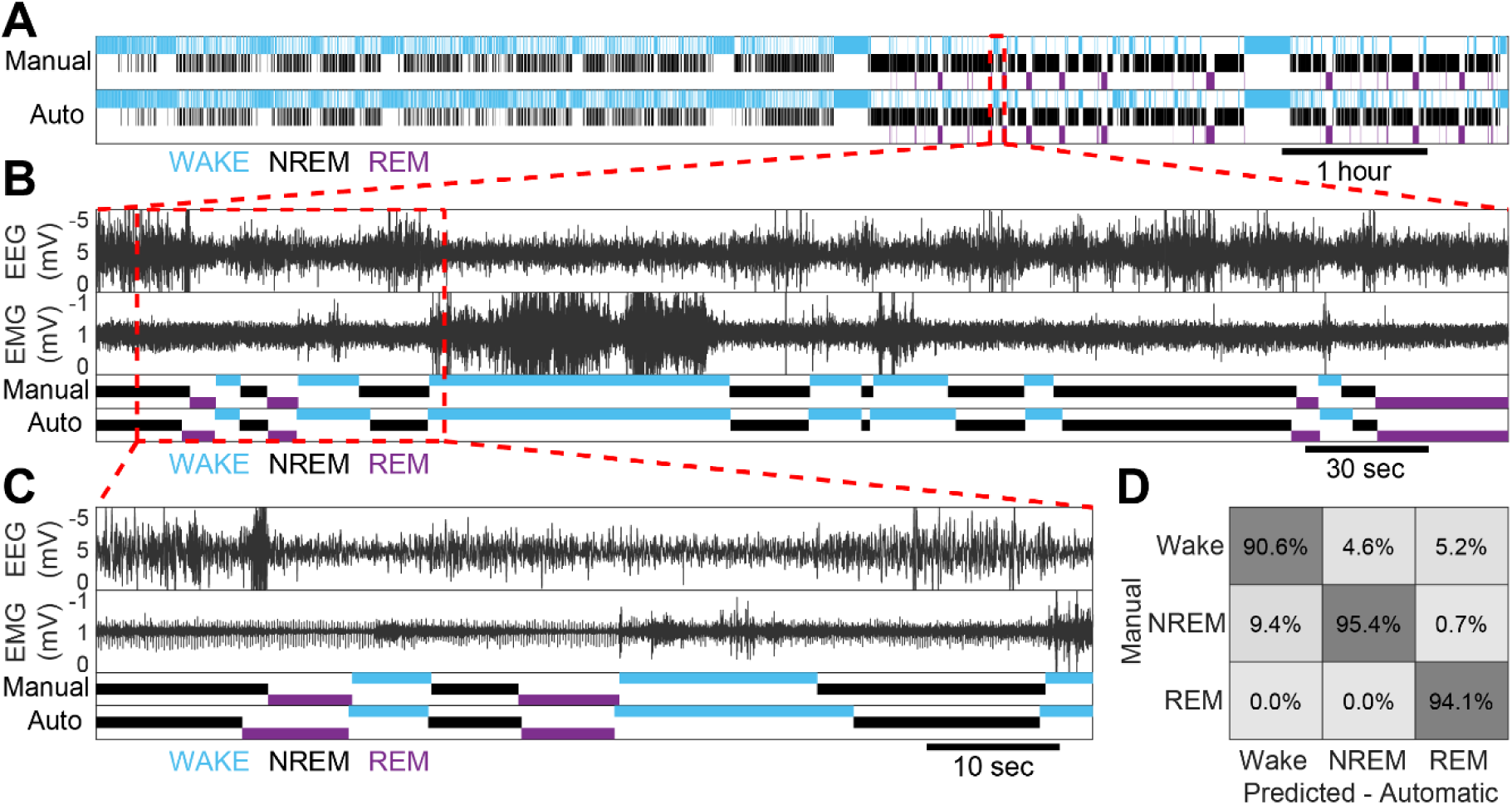
Automated sleep scoring by neural network for extracting falling asleep epochs. (A) Comparison of manual and automated sleep scoring of a representative session ∼9h (top and bottom hypnograms, respectively). Red rectangle marks an example 6-minute segment of vigilance and sleep state changes expanded in panel B. (B) Polysomnography of a representative 6-minute segment. Rows (top to bottom) mark EEG, EMG, and manually and automatically scored vigilance state, as in A. Red rectangle marks further focus on 75 seconds expanded in panel C. (C) Same as in B, but for a shorter 75-second segment. (D) Confusion matrix between the vigilance states scored by automatic and manual sleep scoring over 3 sessions (∼28h). Along the diagonal is the percentage of the automatically scored labels (column) that matched the manually scored labels (row). Outside of the diagonal is the percentage of the automatically scored labels that did not match the manually scored labels but instead the label of the relevant row.

### Falling asleep suppresses sound-evoked post-onset neuronal firing within seconds

To analyze how neural processing and sensory responses in the auditory cortex changed around falling asleep with fine temporal resolution, we defined three intervals around wake-sleep transitions: “wakefulness” (-20 to -5sec before falling asleep), “First seconds of NREM” (+5 to +10sec after falling asleep), and “NREM” (>+60 after falling asleep). Based on our previous study on state-dependent auditory processing (Marmelshtein et al., 2023), we hypothesized that features of auditory cortical processing such as onset responses will maintain their magnitude when falling asleep, whereas features such as post-onset neuronal firing, population synchrony and entrainment to fast click trains (40 clicks/s) may be modulated during the process.

Figure 4A provides an example neuronal response of a representative unit to click stimuli, with trials vertically arranged according to their timing relative to the moment of falling asleep (top to bottom on y-axis). As can be seen for this unit (Figure 4A), as well as the average response (Figure 4B, n=152 units, 7 sessions), the onset response magnitude did not change around falling asleep (comparing wakefulness and first seconds of NREM). In contrast, the post-onset period (30-80 ms following sound onset) showed differences already during the first seconds of NREM sleep, in line with our hypothesis.

**Figure 4.**
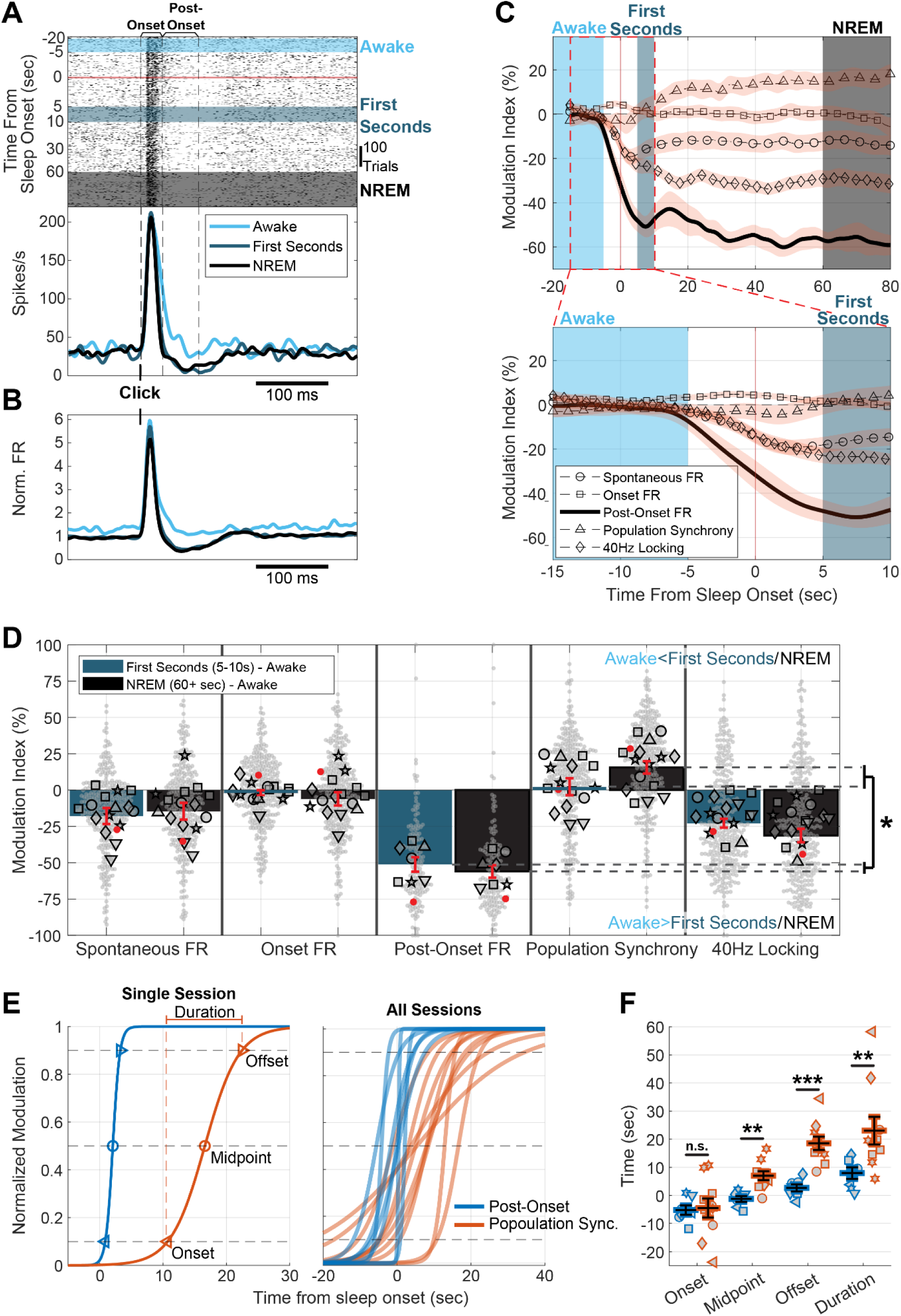
Post-onset firing changes within first seconds of falling asleep. (A) Representative raster and peristimulus time histogram (PSTH) for a unit in response to a click. Raster trials (rows) are ordered according to stimulus time relative to the proximate falling asleep instance. Azure, dark blue, and black shadings and line colors mark if the stimulus occurred while the animal was awake, in the first seconds after falling asleep, or during NREM sleep >60s after falling asleep, respectively. (B) Peristimulus time histogram of the mean normalized response to a click of all responsive units (n = 152, 7 sessions) (C) Top: Mean modulation Index of activity and response features depending on stimulus time relative to the moment of falling asleep for all units (n = 152, 7 sessions for Post-Onset FR; n=233, 15 sessions for 40-Hz locking; and n = 317, 15 sessions, for the other features). Orange shading represents SEM. Bottom: Same as top panel but expanding 25s around the moment of falling asleep. (D) Modulation Index of activity and response features between the states of “first seconds NREM” and “awake” (dark turquoise) and between the states “NREM” and “awake” (black) across units and sessions (n = 152/233/317 units, 7/15 sessions as in C). Bars and error bars represent the mean ± SEM estimated by the Linear Mixed Effects model. Small grey markers represent individual units (red for the representative unit in panel A). Large grey markers represent individual session mean (with each shape representing an individual animal). (E) 4-parameters sigmoid fit of temporal dynamics of post-onset firing (blue) and population synchrony (orange) for a single representative session (left) and all sessions (right). Left pointing triangles, circles, and right pointing triangles mark the “onset”, “midpoint” and “offset” of the sigmoid fit, respectively (10%, 50% and 90% along the slope). Duration is the difference between the offset and onset. (F) Timing of the onset, midpoint and offset relative to the moment of falling asleep, and the duration across sessions for the post-onset sigmoid fit (blue, n=7 sessions) and population synchrony (orange, n=10). *p < 0.05. **p < 0.01. ***p < 0.001.

**E**xpanding the analysis to all units (n=317) and multiple features of auditory processing (Figure 4C,D) revealed that onset response did not significantly change upon falling asleep, i.e., when comparing wakefulness and NREM sleep (Figure 4D, black shading, -6.08% ± 4.65%, p = 0.19, t_316_ = -1.31, LME [Linear Mixed Effects Model]). We then examined the effect of falling asleep on population synchrony, as quantified by “population coupling” (Okun et al., 2015), representing the firing rate correlation of each unit to the local population (Methods). First, we validated that this measure captures the synchrony characteristics of sleep. Indeed, we found that population synchrony is modulated by sleep stages, EEG slow-wave activity (SWA) and sleep pressure (Table S1). Upon Falling asleep, population synchrony, as well as spontaneous firing rate, exhibited moderate modulation (Figure 4C,D, black; 15.6% ± 4.26%, p = 2.9 × 10^-4^, t_316_ = 3.67 for elevation in population synchrony; -14.6% ± 5.81%, p = 0.013, t_316_ = -2.51 for reduction in spontaneous firing, LME). A larger modulation was observed for sustained locking of firing to a 40-Hz click-train (Figure 4C,D, black; -31.3% ± 4.63%, p = 1.2 × 10^-10^, t_232_ = -6.75, LME). The most notable change however, was observed for post-onset neuronal firing (n = 152 units), which was strongly modulated between wakefulness and NREM, as can be seen in Figure 4A-C (black shadings) and as evident quantitatively (-56% ± 4.3%, p = 1.2 × 10^-26^, t_151_ = -13.1, Figure 4D).

We further examined whether changes occur already within the first few seconds after falling asleep or, alternatively, reflect changes due to more consolidated NREM sleep within tens of seconds. As can be seen in the example unit in Figure 4A (dark blue) and the population response (Figure 4B, dark blue), post-onset firing was already significantly modulated within seconds after sleep onset. Quantifying this for the entire dataset (Figure 4C,D, n=317 units) revealed a sharp variation in post-onset firing within seconds of falling asleep (-51.2% ± 4.94%, p = 2.5 × 10^-19^, t_151_ = -10.4, comparing wakefulness to first seconds of NREM). Changes in spontaneous firing rate and 40-Hz Locking, although more moderate, also occurred within seconds (-17.8% ± 5.47%, p = 1.2 × 10^-3^, t_316_ = -3.26, and -22.9% ± 3.02%, p= 8.8e × 10^-13^, t_232_ = -7.57, Respectively). In contrast, changes in population synchrony were slower to occur and did not show significant modulation in the first seconds of NREM (2.26% ± 5.82%, p = 0.69, t_316_ = 0.38). Analysis of variance among the distinct features of cortical auditory processing confirmed that they are differentially modulated within the first second of NREM (p = 2.1×10^-4^, n = 6 animals, Friedman test). In addition, upon entering a deeper state of NREM sleep, processing features were also differentially modulated (p = 1.1×10^-4^, n = 6 animals, Friedman test). Pairwise comparisons revealed that post-onset firing was modulated significantly more strongly than the other four features (p < 3.5×10^-7^, for all four features within first seconds of NREM, and p < 5.8×10^-^8, for all four features within NREM state, df = 151, LME). The slower buildup of population synchrony can be appreciated most clearly when contrasting Figure 4C top panel (robust changes in temporal dynamics at a scale of tens of seconds) with bottom panel (no changes in temporal dynamics in first seconds). Indeed, a direct comparison established that the difference between the first seconds of NREM and deeper NREM was significantly larger for population synchrony modulation than for post-onset modulation (p = 0.023, df = 151, t151 = 2.3, LME).

We sought to quantitively establish the temporal dynamics of post-onset firing and population synchrony without relying on arbitrarily pre-defined intervals (“wakefulness”, “first seconds” and “NREM”). We therefore fitted a 4-parameters sigmoid function to each session average temporal dynamics trace of post-onset firing and population synchrony (Figure 4E, Methods). We used the slope and inflection point (midpoint) sigmoid function parameters to extract information about the timing and duration of changes in each feature, ignoring the magnitude of the modulation (Figure 4E left, 4F). We found that post-onset firing changes were abrupt, with an average duration of 7.9s ± 2.1s across sessions (Figure 4F, n=7 sessions), and coincided precisely with sleep onset (midpoint: -1.3s ± 1s relative to sleep onset, n=7). In contrast, population synchrony changes occurred significantly later (midpoint: 7s ± 1.6s, n=10, p=0.0012, WRST) and gradually materialized (duration: 23s ± 4.9s, p=0.0046, WRST, n_1_=7, n_2_=10). Overall, while post-onset firing changes fully materialized (reaching 90% of sigmoid function asymptote) moments after the animal fell asleep (Figure 4E,F, offset: 2.7s ± 1.2s after sleep onset), population synchrony took significantly longer to fully materialize (offset: 18.5s ± 2.3s, p=1×10^-4^, WRST, n_1_=7, n_2_=10). Spontaneous firing changes were as abrupt as post-onset firing, while 40-Hz locking changes emerged slowly as population synchrony, but tended to occur earlier (Table S2).

Is the large and rapid reduction in post-onset firing a general characteristic of falling asleep, or could it be due to the partial sleep deprivation and high sleep pressure possibly introduced by our behavioral paradigm? To address this issue, we repeated our analysis in Figure 4D, limiting trials to the last two hours of the recovery sleep period (i.e., with weakest sleep pressure). We found that post-onset firing modulation upon falling asleep was virtually identical in magnitude, and occurred within seconds from falling asleep (-48.6% ± 6.8% within first seconds, -55% ± 6.2% for deeper NREM, df=151, LME). Finally, we examined if post-onset firing suppression was associated with intervals of complete neuronal silence (≥50ms in length), i.e., OFF periods. We found that within seconds from falling asleep the probability of silent intervals within the post-onset window ([30,80]ms) increased significantly (p=0.0156, n=7 sessions, WSRT) and to a level comparable to that observed in deeper NREM state (Figure S1).

### Awakening restores sound-evoked post-onset neuronal firing within seconds

Does cortical auditory processing show an opposite profile upon awakening to that observed when falling asleep? To test this, we compared neuronal activity in the seconds just before awakening (“Sleep”, -20 to -5sec before awakening) and just after awakening (“First seconds of wakefulness, +5 to +10sec after awakening). Figure 5A provides a representative example of a unit firing around awakening. As can be seen, post-onset activity rapidly changes from silence during sleep to firing in wakefulness (Figure 5A,B), whereas population synchrony is minimally affected at these time-scales (Figure 5C). Quantifying these effects across the entire dataset (n=315 units, Figure 5D), we found a modest albeit significant increase in onset firing rate that occurred in the first seconds of wakefulness (11.7% ± 3.82%, p = 2.6 × 10^-3^, t_314_ = 3.04, LME). Similarly, we found moderately higher modulation indices representing elevation in spontaneous firing rate (19.8% ± 6.72%, p = 3.5 × 10^-3^, t_314_ = 2.94) and 40-Hz locking (21.4% ± 3.28%, p = 9.7 × 10^-10^, t_156_ = 6.51). As observed upon falling asleep, changes in post-onset firing were most pronounced and occurred within the first seconds of state transitions (n = 145 units, 42.3% ± 5.46%, p = 1.6 × 10^-12^, t_144_ = 7.74). Population synchrony, as was the case when falling asleep, did not significantly change within the first few seconds around state transitions (-5.2% ± 4.2%, p = 0.22, t_314_ = -1.23). Together, awakening exhibits a “mirror-image” of the changes in cortical auditory activity to those found around falling asleep, with post-onset firing remaining the aspect that changes most strongly and rapidly. Analysis of variance among the distinct features of cortical auditory processing confirmed that they are differentially modulated within the first second of wakefulness (p = 3.3×10^-4^, n = 6 animals, Friedman test). Pairwise comparisons revealed that post-onset firing was modulated significantly more strongly than the other four features (p < 3.8×10^-6^, df = 144, for all four features, LME).

**Figure 5.**
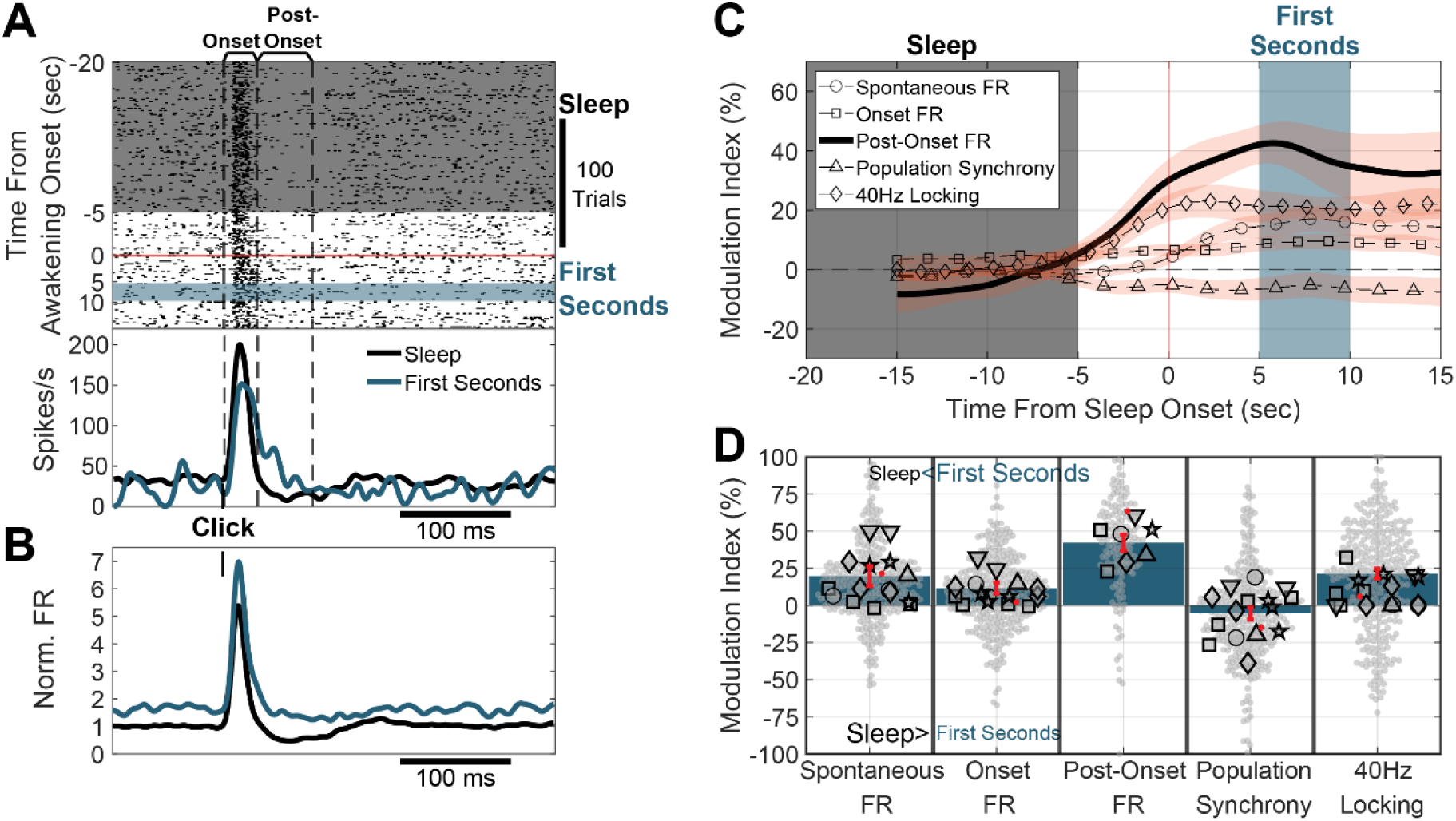
post-onset firing changes within first seconds of awakening. (A) Representative raster and peristimulus time histogram (PSTH) for a unit in response to a click. Raster trials (rows) are ordered according to the trial’s stimulus time relative to the proximate awakening instance. Black and blue shadings mark if the stimulus occurred while the animal was asleep (NREM) or in the first seconds after waking up, respectively. (B) Peristimulus time histogram of the mean normalized response to a click of all responsive units (n = 145, 7 sessions) during sleep and first seconds of wakefulness. (C) Mean modulation Index of activity and response features depending on stimulus time relative to the moment of awakening. Orange shading represents the SEM. (D) Modulation Index of activity and response features between “first seconds wakefulness” and “sleep” states across units and sessions (n = 145, 7 sessions for Post-Onset FR; n = 157, 15 sessions for 40-Hz locking; and n = 315, 15 sessions for the other features). Bars and error bars represent the mean±SEM estimated by the Linear Mixed Effects model. Small grey markers represent individual units (red for the representative unit in panel A). Large grey markers represent individual session mean (with each shape representing an individual animal).

### Falling asleep modulates EEG auditory-evoked potentials within seconds

Are changes in neuronal post-onset firing and population synchrony also captured by the EEG? We sought to test this, seeking to relate our findings to the rich literature studying falling asleep with EEG. To this end, we followed the same approach, comparing the EEG recorded from a screw over the frontal lobe across three predefined intervals (Fig 6A-B) of “wakefulness” (-20 to -5sec before falling asleep), “First seconds of NREM” (+5 to +10sec after falling asleep), and “NREM” (>+60 after falling asleep). EEG auditory evoked potentials (AEPs) in response to click stimuli exhibited rapid changes within seconds upon falling asleep (Fig 6A). As was the case in auditory cortex neuronal activity, an initial (30ms) large positive deflection in the AEP was unchanged before and after sleep onset. By contrast, a later (50-140ms) large negative deflection in the AEP occurring at the time of the neuronal post-onset firing period was strongly modulated within the first seconds of NREM sleep (magnitude of 301±64% relative to wakefulness, p=0.016, n=7 sessions, Wilcoxon Sign-Rank Test [WSRT]). It was then followed, but only after sleep onset, by an even later (180-280ms) positive deflection and power increase in the spindle frequency range (Figure S2), suggesting similarity to an evoked K-complex.

**Figure 6.**
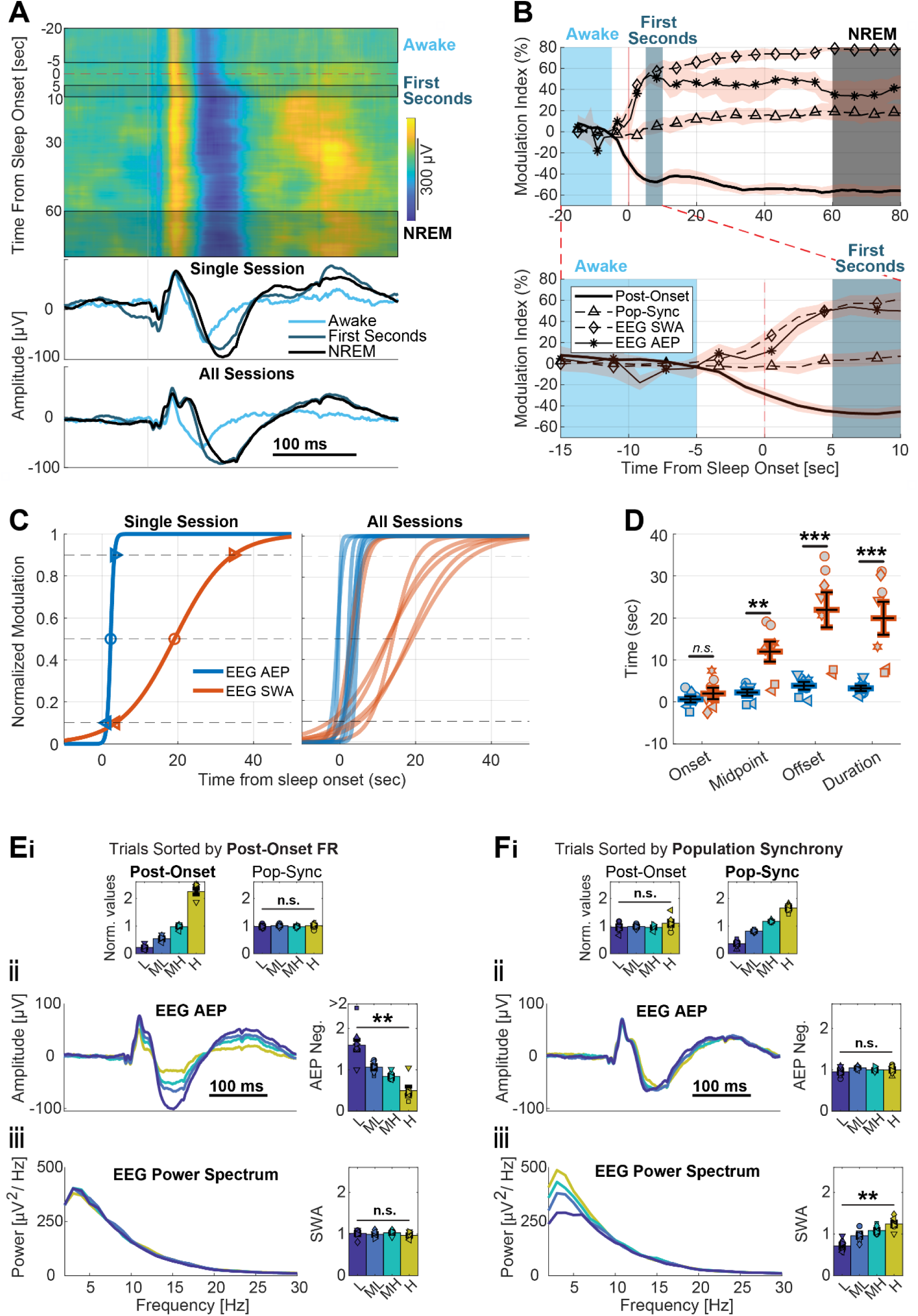
EEG AEP changes within first seconds of falling asleep. (A) Top: changes upon falling asleep in frontal EEG auditory evoked potential (AEP) following a click stimulus in a single representative session. Individual trials were sorted according to stimulus time relative to the proximate falling asleep instance. Each row represents an average AEP of 100 consecutive trials. Azure, dark blue and black shadings mark if stimuli occurred while the animal was awake, in the first seconds after falling asleep, or during NREM sleep >60s after falling asleep, respectively. Middle: Average AEPs for the three different intervals in a single session. Bottom: Same as middle panel, but averaged across all sessions (n=7). (B) Top: Temporal dynamics upon falling asleep of the modulation index of frontal EEG features, as well as auditory cortex neuronal features for comparison, averaged across all sessions (n=7). Frontal EEG AEP negative deflection and slow-wave activity (SWA) are marked by asterisk and diamond traces, respectively. Post-onset firing and population synchrony in auditory cortex neurons are marked by thick and triangle (dashed) traces, respectively. Bottom: same as top panel but with temporal axis expanded. (C) 4-parameters sigmoid fit of temporal dynamics of EEG AEP negative deflection (blue) and slow-wave activity (SWA, orange) for a single representative session (left) and all sessions (right). Left pointing triangles, circles, and right pointing triangles mark the “onset”, “midpoint” and “offset” of the sigmoid slope, respectively (10%, 50% and 90% along the slope). Duration is the difference between the offset and onset. (D) Timing of the onset, midpoint and offset relative to the moment of falling asleep, and the duration across all sessions (n=7) for the AEP negative deflection sigmoid fit (blue) and SWA (orange). (E) trials were sorted into four quartiles based on their relative auditory cortex post-onset firing (low post-onset FR (L), medium-low (ML), medium-high (MH) and high(H). (Ei) Auditory cortex normalized post-onset firing (left) and population synchrony (right) averaged across all sessions for the four different trial groups. (Eii) Average frontal EEG AEP (left) and its negative deflection magnitude (right) for the four different trials groups across all sessions. (Eiii) Average baseline (pre-stimulus) power spectrum (left) and slow-wave activity (right, 0.5-4 Hz power) for the four different trials groups across all sessions. (F) same as E but with trials sorted into four quartiles based on their relative population synchrony in auditory cortex (low population synchrony (L), medium-low (ML), medium-high (MH) and high(H)). **p < 0.01. ***p < 0.001.

We quantified the temporal dynamics of this effect across all sessions by obtaining the AEP negative deflection magnitude (methods) and calculated a running modulation index (Figure 6B, trace with asterisks) as for the previous auditory cortex neuronal response features (Figure 4C). The AEP negative deflection showed temporal dynamics qualitatively similar to the post-onset response in the auditory cortex, both reaching their maximal change within seconds after falling asleep and then plateauing. Qualitatively similar results, but with opposite direction, were observed upon awakenings (Figure 7). We also examined how EEG baseline (pre-stimulus) slow wave activity (SWA, power 0.5-4 Hz) depends on falling asleep (Figure 6B, diamonds), given its relation to synchronous neuronal activity (Vyazovskiy et al., 2009). SWA increased within seconds upon falling asleep (Figure 6B, 277±40% relative towakefulness, p=0.016, n=7 sessions, WSRT) but also continued to increase slowly afterwards (Figure 6B, 437±56% relative to wakefulness), exhibiting a significant further increase from the “First seconds of NREM” to subsequent “NREM” intervals (p = 0.031, n=7 sessions, WSRT).

**Figure 7.**
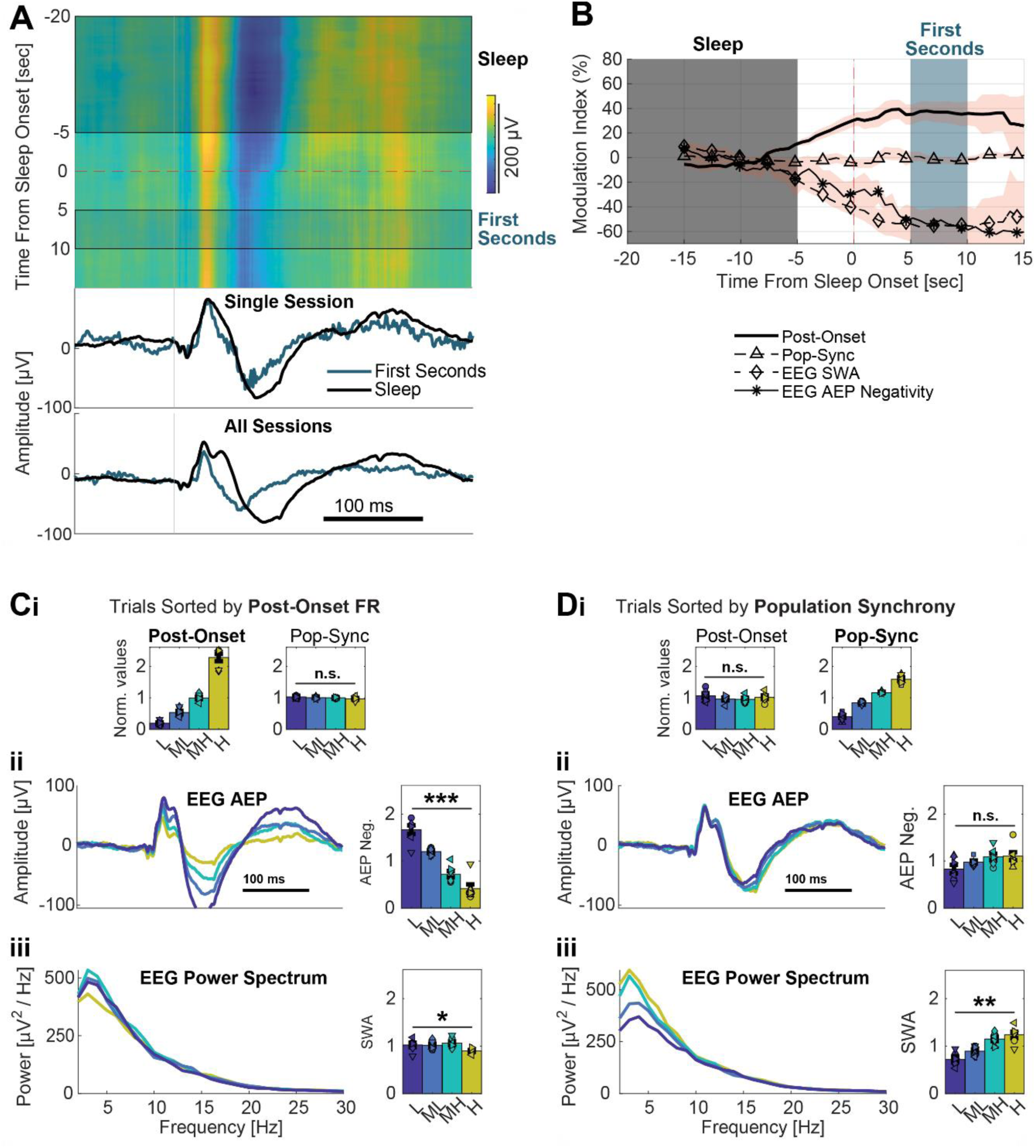
EEG AEP changes within first seconds of awakening. Same as Figure 6 but for awakenings. (A) Top: changes upon awakenings in frontal EEG auditory evoked potential (AEP) following a click in a single representative session. Individual trials were sorted according to stimulus time relative to the proximate awakening instance. Each row represents an average AEP of 100 consecutive trials. Black and blue shadings mark if the stimulus occurred while the animal was asleep, or during the first seconds after waking up. Middle: Average AEPs for the two different intervals in a single session. Bottom: Same as middle panel, but averaged across all sessions (N=7). (B) Temporal dynamics upon waking up of the modulation index of frontal EEG features, as well as auditory cortex neuronal features for comparison, averaged across all sessions (N=7). Frontal EEG AEP negative deflection and slow-wave activity (SWA) are marked by asterisk and diamond traces, respectively. Post-onset firing and population synchrony in auditory cortex neurons are marked by thick and triangle (dashed) traces, respectively. (C) trials were sorted into four quartiles based on their relative auditory cortex post-onset firing (low post-onset FR (L), medium-low (ML), medium-high (MH) and high(H). (Ci) Auditory cortex normalized post-onset firing (left) and population synchrony (right) averaged across all sessions for the four different trial groups. (Cii) Average frontal EEG AEP (left) and its negative deflection magnitude (right) for the four different trials groups across all sessions. (Ciii) Average baseline (pre-stimulus) power spectrum (left) and slow-wave activity (right, 0.5-4 Hz power) for the four different trials groups across all sessions. (D) same as C but with trials sorted into four quartiles based on their relative population synchrony in auditory cortex (low population synchrony (L), medium-low (ML), medium-high (MH) and high(H)). *p < 0.05. **p < 0.01. ***p < 0.001.

We sought to capture the temporal dynamics of these phenomena without relying on arbitrarily defined intervals. Fitting again a 4-parameters sigmoid function to each session data (Figure 6C), we found that changes in the AEP negative deflection emerged instantly (Figure 6C,D, duration: 3.2s ± 0.65s, n=7 sessions) and coincided with sleep onset (midpoint: 2.2s ± 0.76s following sleep onset, n=7). This was in stark contrast to changes in EEG SWA that emerged significantly later (midpoint: 12s ± 2.4s, p =0.007, WRST, n_1_=7, n_2_=7) and more gradually (duration: 19.9s ± 3.9s, p=5.8×10^-4^, WRST, n_1_=7, n_2_=7). Furthermore, changes in the AEP negative deflection fully materialized moments after the animal fell asleep (Figure 6C,D, offset: 3.8s ± 0.9s after sleep onset), where EEG SWA took much longer (offset: 22s ± 4.2s, p=5.8×10^-4^, WRST, n_1_=7, n_2_=7), closely reproducing the timing of the auditory cortex neuronal measures, post-onset firing (offset: 2.7s ± 1.2s) and population synchrony (offset: 18.5s ± 2.3s), respectively.

Next, we examined the relation between neuronal activity measures (post-onset firing and population synchrony) and EEG AEP measures (AEP negative deflection and SWA) by testing their association at the single-trial level (Figure 6E). To go beyond their shared temporal dynamics, we selected groups of trials with similar timing relative to the moment of falling asleep and divided them into four quartiles according to their auditory cortex post-onset firing (methods). This created four groups of trials with different average post-onset firing (Figure 6Ei left) that were temporally balanced across the falling asleep process (i.e., similar number of trials before, during, and after falling asleep in each group). The four trial groups did not differ in their average auditory cortex population synchrony (Figure 6Ei right, p=0.543, n=7 sessions, Friedman test) implying that the two neuronal measures are governed by fundamentally different processes. Strikingly, when extracting the EEG AEP for each of these four groups, strong differences emerged (Figure 6Eii left) with significant and reliable parametric modulation of the negative deflection magnitude (Figure 6Eii right, p=0.0083, n=7 sessions, Friedman test). Importantly, no changes in baseline SWA were observed between the four different groups (Figure 6Eiii, p=0.419, n=7 sessions, Friedman test). Thus, post-onset firing in auditory cortex is specifically associated with the AEP negative deflection.

We then performed the same analysis, but now dividing trials based on their relative auditory cortex population synchrony (Figure 6F). This process now created four groups that were different in their population synchrony (Figure 6Fi right), but that showed no difference in their average post-onset firing (Figure 6Fi left, p=0.319, n=7 sessions, Friedman test). This led to an exactly opposite pattern, with no observable EEG AEP differences between the groups (Figure 6Fii, p=0.419, n=7 sessions, Friedman test) but with significant and reliable parametric differences in SWA (Figure 6Fiii, p=0.002, n=7 sessions, Friedman test). Qualitatively similar results were also observed upon awakenings (Figure 7C-D). Importantly, dividing the trials based on other neuronal features, such as the relative spontaneous firing (Figure S3A), or onset firing (Figure S3B), revealed no association with either EEG measure. Overall, this analysis establishes that reduced post-onset firing in auditory cortex neurons is selectively associated with the EEG AEP late negative deflection, while auditory cortex neuronal population synchrony is selectively associated with EEG slow wave activity, thereby revealing a robust double-dissociation between the two auditory cortex measures and the two EEG phenomena.

## Discussion

We studied how neuronal spiking activity and responses to clicks in early rat auditory cortex change in the seconds around falling asleep and waking up. We found that increased post-onset neuronal silent periods (OFF states) and moderate reductions in spontaneous firing rates - both known to occur in NREM sleep (Marmelshtein et al., 2023) - changed abruptly around the first seconds of falling asleep. These changes in activity and responsivity during the first moments of sleep reached their full magnitude as observed in later consolidated NREM sleep (<60 sec from the moment of falling asleep). By contrast, an increase in neuronal population synchrony only changed substantially during later (>60 sec) NREM sleep without robust modulations around the seconds of falling asleep. Onset response magnitude was unaffected by falling asleep or awakening, echoing previous studies showing its invariance during sleep (Issa and Wang, 2008; Nir et al., 2013). During the first seconds after awakenings, we found opposite changes in neuronal activity and responsivity (rapid and robust decrease in the occurrence of post-onset silent periods, and rapid moderate elevation in spontaneous firing rates). These changes showed identical temporal dynamics, but with opposite directionality, to those observed when falling asleep. Changes in EEG activities including modulations in baseline SWA and the AEP were also observed upon falling asleep and awakening. SWA was selectively associated with changes in auditory cortex population synchrony, whereas the AEP negative deflection was selectively associated with post-onset neuronal silence. Together, the results indicate that some changes in auditory cortex activity occur within seconds around wake-sleep transitions whereas specific aspects such as population synchrony vary with slower dynamics. Auditory-induced silent OFF periods emerge as the aspect that changes most robustly around state-transitions, showing abrupt and large changes in its magnitude within seconds. The tight correlation between auditory cortex activity and the EEG recorded over the frontal lobe imply that these changes are not limited to the auditory cortex.

How do our findings relate to previous studies on auditory responses across different arousal states? A recent study on the effects of gradual propofol anesthesia reported that auditory processing changes occurred slowly, closely tracking increasing anesthetic concentration rather than abrupt loss of behavioral responsiveness (Bergman et al., 2022). Similarly, we previously demonstrated that post-onset neuronal firing decreases gradually with rising sleep pressure over hours of sleep deprivation (Marmelshtein et al., 2023). These studies indicate that gradual changes in arousal often lead to gradual adjustments in auditory processing. In contrast, our current work uniquely examines rapid transitions between wakefulness and sleep, revealing that neural responses can shift sharply within seconds — evident in the sudden decrease in post-onset firing at sleep onset. We propose that the dynamics of arousal-related neural changes depend on the nature of the state transition: gradual changes in arousal (e.g., mounting sleep pressure or deepening anesthesia) yield gradual neural adjustments, while abrupt state changes (e.g., falling asleep or awakening) result in sharp neural shifts.

Beyond neuronal correlates in specific pathways such as the auditory cortex, what do our findings reveal more generally about falling asleep – is it fundamentally a gradual or abrupt process? While extensive evidence shows that many electrical, physiological and behavioral phenomena change gradually in the span of minutes (Ogilvie, 2001; Eban-Rothschild et al., 2016; Lacaux et al., 2023), some EEG phenomena and specific measures of behavioral responsiveness do show fast temporal dynamics (Ogilvie, 2001; Strauss et al., 2022). We find evidence supporting both of these seemingly contradicting themes. On the one hand, post-onset neuronal silence changes rapidly within just few seconds upon sleep onset. Exposing these fast dynamics relied critically on our ability to sleep score our data at high temporal resolution (1 second). On the other hand, population synchrony displayed different temporal dynamics, developing gradually over a span of 20-30 seconds. Similarly, measures of EEG information and connectivity also change on a slow time-scale (Strauss et al., 2022). This implies that these two measures may reflect different underlying neural processes both accompanying sleep onset, one that is continuous and gradual, and another abrupt.

Examining the effects of falling asleep (and awakening) on EEG activity helps connect the observations at the neuronal level to the non-invasive literature. The change in baseline (pre-stimulus) EEG SWA upon falling asleep is not necessarily surprising, as these changes comprise the criteria used to define wake-sleep transitions. By contrast, the fast change in the K complex-like AEP negative deflection reached maximal modulation within seconds of falling asleep or awakening, thereby providing independent evidence for rapid changes in brain activity upon sleep state transitions. Investigating trial-by-trial variability allowed to establish direct link between EEG and neuronal-level phenomena, revealing a double-dissociation between the EEG AEP and post-onset neuronal silence (changing rapidly and reaching maximum change within seconds) versus population synchrony and ongoing EEG SWA (continuing to change over longer time scales after state transitions).

The functional significance of increase in post-onset neuronal silence remains unclear. Considering the strong association with the EEG AEP, it is highly likely that neuronal silence as captured in extracellular recordings reflects a state of neuronal bistability where neurons alternative between active (up) and hyperpolarized silent (down) states, a key feature of sleep and unresponsive states (Massimini et al., 2007; Vyazovskiy et al., 2011, 2013; Marmelshtein et al., 2023). While such bistability does not interfere with short-lived localized onset responses, it can disrupt complex neural processing by breaking-off sustained and widespread propagation of sensory signals (Massimini et al., 2005; Pigorini et al., 2015). Indeed, states of drowsiness and sleep associated with increased bistability affect processing requiring integration beyond local activity patterns. Accordingly, drowsiness preferentially disrupts perceptual discrimination, rather than perceptual detection (Xu et al., 2023). Moreover, measures of complex neural processing, such as predictive coding, are preferentially affected by sleep onset (Strauss et al., 2015), and baseline complexity measures were maximally attenuated in NREM sleep when stimuli evoked a k-complex, reflecting high neuronal bistability (Andrillon et al., 2016).

Certain limitations in our study require explicit acknowledgment. First, it is not trivial to generalize results about the precise dynamics around state transitions from rodents to humans and other species. Since rodents exhibit more fragmented sleep and shorter bouts of sleep and wakefulness (Lo et al., 2004), the first few seconds of sleep or wakefulness in rats may or may not be directly comparable to the first few seconds of longer sleep bouts. Nevertheless, the fact that some aspects of auditory processing changed robustly and abruptly (post-onset firing) whereas other aspects changed more modestly (ongoing firing rates) or more gradually (population synchrony) goes beyond differences across species. Second, while our recordings focused on (early) auditory cortex and AEPs in the EEG, the spatial scale of these effects remains unclear. Further studies are needed to examine to what extent such findings can be generalized to other sensory modalities and cortical areas. However, the selective relationship between auditory cortex post-onset responses and the frontal EEG AEP negative deflection suggests that such effects may be widespread. Third, by focusing on a limited set of simple auditory stimuli (click trains), we achieved frequent repetitions and high temporal resolution during sleep onset and awakening. However, whether sleep transition affects the encoding or processing of complex, behaviorally relevant sounds remains an open question. Finally, some limitations relate to our extracellular recordings, discussed also previously (Marmelshtein et al., 2023); these include a bias for larger pyramidal cells with high baseline firing rates as well as lack of information on cortical layer and cell types, possibly biasing our results towards specific neuronal subpopulations.

In conclusion, our investigation reveals the temporal dynamics and magnitude of modulation in different features of auditory processing upon falling asleep and awakening. Our findings extend previous studies in showing that specific neural signatures of auditory responses, particularly the degree of post-onset firing, are robustly influenced by vigilance states and their transitions (Marmelshtein et al., 2023). A rapid and robust dramatic increase in post-onset neuronal silence, occurring within seconds of falling asleep, contrasts with the slower modulation observed in population synchrony, and emerges as a robust signature of altered responsivity of the sleeping brain to external sensory stimuli.

## Methods

### Animals

The study involved electrophysiology experiments in six adult male Wistar rats (weight: 300-350g, age: >12 weeks). Rats were kept in transparent Perspex cages with food and water available ad libitum. The ambient temperature was maintained between 20-24 Celsius, and there was a 12:12 hours light/dark cycle with light onset at 10:00 AM. All experimental procedures, including animal handling, wheel cycle paradigm, and surgery, were conducted in accordance with the National Institutes of Health’s Guide for the care and use of laboratory animals and approved by the Institutional Animal Care and Use Committee of Tel Aviv University.

### Surgery and electrode implantation

Before surgery, DiI fluorescent dye (DiIC18, Invitrogen) was applied to microwire arrays under microscopic control to aid in localization. Generally, surgical procedures followed our previous protocols (Sela et al., 2020). First, isoflurane (4%) was used to induce general anesthesia, and animals were then placed in a stereotactic frame (David Kopf Instruments). During the rest of the surgery, anesthesia (isoflurane, 1.5-2%) and a 37C body temperature (closed-loop heating pad system, Harvard Apparatus) were maintained. Animals were given antibiotics (Cefazolin, 20 mg/kg i.m.), analgesia (Carpofen, 5 mg/kg i.p.), and dexamethasone (0.5 mg/kg, i.p.). Their scalp was shaved, and liquid gel (Viscotears) was applied to protect their eyes. Subcutaneous infusion of lignocaine (7 mg/kg) was performed before incision, and then the skull was exposed and cleaned. Two frontal screws (one on each hemisphere, 1mm in diameter) and a single parietal screw (left hemisphere) were placed in the skull for recording EEG. Two screws were placed above the cerebellum, serving as reference and ground. Two single-stranded stainless-steel wires were inserted to the neck muscles to record EMG. EEG and EMG wires were soldered to a head-stage connector (Omnetics). Dental cement was applied to cover all screws and wires. A small craniotomy was performed over the right hemisphere, and the dura was meticulously dissected. An auditory cortex-targeting 16-electrode microwire array (Tucker-Davis Technologies, TDT, 33 or 50mm wire diameter, 6-6.5 mm long, 15 tip angle; arrays consisting of 2 rows of 8 wires, with 375mm medial-lateral separation between rows and 250mm anterior–posterior separation within each row) was implanted diagonally (angle of 28 degrees, see Figure 1B) with insertion point center coordinates of P: -4.30mm, L: 4.5mm relative to Bregma, and inserted until a final depth of 4.6mm was reached. Silicone gel (Kwik-Sil; World Precision Instruments) was applied to cover the craniotomy, and Fusio (Pentron) was used to fix the microwire array in place. After surgery, chloramphenicol 3% ointment was applied topically, and buprenorphine (0.025 mg/kg s.c.) was given systemically for additional post-operative analgesia. Dexamethasone (1.3 mg/kg) was given with food in the days following the surgery to reduce pain and inflammation around implantation.

### Histology

After conducting the experiments, we confirmed the position of electrodes in 4 out of 6 animals through histology, as shown in Figure1B. Animals were anesthetized with 5% isoflurane and perfused with 4% paraformaldehyde. The brains were refrigerated in paraformaldehyde for a week and then cut into 50-60mm serial coronal sections using a vibrating microtome (Leica Biosystems). The sections were stained with fluorescent cresyl violet/Nissl (Rhenium). The histological verification ensured that the electrodes were located within areas Au1/AAF/AuD as defined by Paxinos and Watson (Paxinos and Watson, 2006).

### Electrophysiology

Data were acquired using a RZ2 processor (TDT). Microwire extracellular activity was digitally sampled at 24.4 kHz (PZ2 amplifier), while EEG and EMG were pre-amplified (RA16LI, TDT) and sampled at 256.9 Hz (PZ2 amplifier, TDT). Spike sorting was performed with “wave_clus” (Quiroga et al., 2004), which utilized a detection threshold of five SDs and automatic superparamagnetic clustering of wavelet coefficients. The clusters were then manually selected, refined, and tagged as either multi-units (MU) or single-units (SU) based on their stability throughout the recording, quality of separation from other clusters, consistency of spike waveforms, and inter-spike interval distributions (N_MU_ = 332, N_SU_ = 88), as described in (Nir et al., 2015). We verified the accurate localization of the auditory cortex, which includes A1, the anterior auditory field, or the dorsal auditory cortex, using histology (Figure 1B) and by analyzing the units’ response latency to auditory stimuli (Marmelshtein et al., 2023).

### Experimental Design

Following surgery, the subjects were housed separately and introduced to the motorized running wheel for a few hours every day for habituation (Figure 1A). They were then gradually introduced to the periodic wheel rotations to be performed in the first experimental phase (Figure 1D, see below). We conducted 16 experimental sessions in the following manner. At the start of the day (10 AM), the rats were moved from their home cage to a motorized running wheel (Figure 1A, Model 80860B, Lafayette Instrument) placed within a sound-attenuation chamber (-55dB, H.N.A) and underwent five hours of periodic wheel rotation. During the first experimental phase, the wheel was rotated for 3 seconds every 36-39 seconds, which constituted intervals when rats could behave freely (run, groom, or fall asleep). In the second experimental phase, rats were allowed to rest undisturbed for an additional 5 hours in the (fixed) wheel, which constituted a recovery sleep opportunity (Figure 1D).

The rats were kept in the wheel apparatus under the ultrasonic speaker for the entire session. Auditory stimulation was delivered intermittently throughout both the sleep deprivation and recovery sleep periods, irrespective of the wheel’s movement.

### Auditory stimulation

We used Matlab (MathWorks) to compile sound waveforms that were then transduced into voltage signals using a high-sampling rate sound card (192 kHz, LynxTWO, Lynx). These signals were amplified (SA1, TDT) and played through a magnetic speaker (MF1, TDT), mounted 60 cm above the motorized running wheel. We conducted two different auditory experiments over separate sessions/days:

### Auditory paradigm A

This paradigm was used in 7 sessions in 6 animals: Stimuli included click trains and a set of Dynamic Random Chords (DRC)(Linden et al., 2003). Click trains lasted 500ms and were played at rates of either {2, 10, 20, 30, 40} clicks/sec (Figure 1C). A typical 10-hour session contained 2000 blocks, each consisting of a single DRC stimulus and a single repetition of each click train (presented at random order), with an inter-stimulus interval of 2s and ±0.25s jitter.

### Auditory paradigm B

This paradigm was used in 9 sessions in 5 animals: The stimuli used in the study included a 40 clicks/second click-train and a set of DRC stimuli. Each 10-hour session had 600 blocks, with each block consisting of a single DRC stimulus and 4 repetitions of the 40 Hz click train, presented in random order. The inter-stimulus interval was 2 seconds, with a ±0.25-second jitter. Both paradigms included an 8-second inter-stimulus interval every 2 minutes.

### Acoustic sound-level measurements

We used a high-quality ultrasonic microphone (model 378C01, PCB Piezotronics) to measure sound levels at 12 positions around the circumference of the wheel apparatus (anterior-posterior axis) and at five positions along its width (left-right axis) with a spacing of 2cm. This covered the area where animals were present during the experimental sessions. We found that the root-mean-square (RMS) intensity in response to 2-second white noise sounds had minimal variance at different positions along the anterior-posterior axis, with a mean absolute deviation of 0.7±0.3dB (±Std. Dev) and a maximum difference of 2.4dB between any two positions. Similarly, along the left-right axis, we recorded a mean absolute difference of 0.4±0.3dB (±Std. Dev.) and a maximum difference of 1.5dB. When we examined sound levels at different frequency bands (1, 2, 4, 8, 16, 32, and 64 KHz, in response to the same white noise stimulus) at different positions, we found similarly low variation (all below 4dB)).

### Sleep scoring and identification of state transitions

In each session (n=16), at least one hour (the last hour, representing recovery sleep opportunity period) was manually sleep scored, employing visual inspection of EEGs, EMGs, and video/behavior as in previous studies (Vyazovskiy et al., 2011; Rodriguez et al., 2016; Sela et al., 2020). In an additional three sessions, manual sleep scoring of the entire 8-9h experiment was performed and used for assessing the performance of automatic sleep scoring (Figure 3).

Before any sleep scoring, we excluded any periods when the wheel was moving (forced running bouts during the first experimental phase). Next, we categorized different time periods to either active-wakefulness with behavioral activity (e.g., locomotion, grooming) as confirmed by video, quiescent-wakefulness (low-voltage high-frequency EEG activity and high tonic EMG with occasional phasic activity), NREM sleep (high-amplitude slow wave activity and low tonic EMG activity), REM sleep (low-amplitude wake-like frontal EEG co-occurring with theta activity in parietal EEG and flat EMG, Figure 3), or unknown periods not analyzed further (e.g. state transitions, to conservatively avoid these epochs in subsequent analysis).

Next, the 1h+ manually-scored data in each session was used to train an automated sleep-scoring algorithm using convolutional neural networks (CNNs) (Barger et al., 2019). The algorithm was fed with the manual sleep scoring vector along with the frontal EEG signal (low-pass filtered < 60Hz), and the EMG signal (band-pass filtered 10Hz – 125Hz). Manually scored labels, originally tagged in 1ms resolution, were re-labeled into 1sec epochs, where each epoch was labeled according to the most frequent manual label in that interval. For each rat, a personalized CNN was trained using the manually labeled data of all the sessions of the same rat (1-4 sessions per rat). Weights for mixture z-scoring, for both training and classification, were calculated using the relative prevalence of each state in the data used for training. Epoch length was 1 sec. The CNNs trained with consideration of 9 epochs surrounding each epoch scored (four epochs preceding and four epochs following the current epoch). Calibration data for mixture z-scoring for classification was also provided as the training data. Using the relevant rat’s CNN for each session, the entire data set was classified into the sleep scoring labels. After applying the classification method, we designed a post-processing method for the labels to meet our goal for early detection of true fall-asleep (awakening) transitions. In which case, for each session, the epochs labels were examined from the last one to the first one. Epochs with low certainty levels in all states, i.e., below 50%, were set to be the same as the label of the following epoch (the later in time). Then, epochs that were labeled differently from their two neighboring (preceding and following) epochs, and their neighbors had the same label, were changed to match their neighbors.

This occurred rarely, in only 1.4% ± 0.6% of 1s epochs (mean ± std, n=15 sessions). Any epochs containing times when the wheel was moving were excluded from the analysis.

Extracting falling asleep and awakening episodes, based on the automatic sleep scoring, was performed as follows. We considered a falling asleep instance as a continuous sequence (≥2 sec) of NREM epochs (epoch = 1 sec), preceded by a continuous sequence (≥2 sec) of quiescent-wakefulness epochs. The beginning of the first NREM epoch was considered, for our matter, the moment of falling asleep. For the subsequent analysis of neural activity, a symmetrical time window around the falling asleep moment was taken. The window extended as long as possible without including periods of rat movement, wheel movement, or epochs that are not part of quiescent-wakefulness or NREM sequences. Falling asleep instances were discarded from the subsequent analysis if more than one falling asleep transition occurred within the same 36-39sec interval between consecutive wheel cycles. Awakening instances were extracted only from the spontaneous sleep opportunity phase. We considered an awakening instance in a similar way, a continuous sequence of quiescent-wakefulness epochs (≥2 sec), preceded by a continuous sequence (≥2 sec) of NREM epochs. The beginning of the first quiescent-wakefulness epoch was considered, for our matter, the moment of awakening. For the subsequent analysis of neural activity, a symmetrical time window around the awakening moment was taken. The window extended as long as possible without including periods of rat movement, or epochs that are not part of quiescent-wakefulness or NREM sequences. Falling asleep and awakening instances with a time window shorter than 3 sec were not analyzed. Across the three fully manually scored sessions, automatic and manual scoring agreed on the occurrence of falling-asleep instances in 82.4% ± 1% of inter-wheel-rotation intervals (first experimental phase). In these cases, sleep onset timings were closely aligned, with mean ± standard deviation time differences of -1.5 ± 4.6 s, 0.1 ± 5 s, and 2.8 ± 4.4 s between automatic and manual scoring across the three sessions.

## QUANTIFICATION AND STATISTICAL ANALYSIS

### Analysis of auditory responses across states

Auditory stimuli presented during the times of wheel movement or while the rat was active (locomotion, grooming for example) were excluded from analysis.

### Analysis of neuronal responses to click trains (Figures 4C-D, 5C-D)

To measure spontaneous firing rate, we calculated the average firing rate during the [-500,0]ms window leading up to the click-train stimuli. For post-onset firing rate, we calculated the average firing rate during the [30,80]ms window. We analyzed onset response to different click rates using smoothed peri-stimulus time histograms (PSTH) with a Gaussian kernel (σ=2ms). The onset response was determined by extracting the highest firing rate during the [0,50]ms window of the smoothed PSTH. Firing rate locking to 40 Hz click trains was obtained by calculating the mean firing rate for each phase (in 1ms bins) during the inter-click intervals in the [130,530]ms window. Then, firing rate locking was defined by the minimum firing rate (during the least preferred phase relative to the click) subtracted from the maximum firing rate (during the most preferred phase). Population synchrony was defined as population coupling (Okun et al., 2015), calculated as the Pearson correlation coefficient of each unit’s firing rate within 50ms bins to the average firing rate of the neuronal population, during a single baseline period ([-1000,0]ms, constituting 20 consecutive bins of 50ms). It thus captures the low-frequency correlation (≤20Hz, due to the 50ms bins) of a unit firing rate to that of the entire neuronal population, within a single trial baseline period. We calculated population coupling for each trial baseline period and then averaged for all trials in a given condition.

To assess whether population synchrony accurately reflects the synchrony characteristics of sleep (Table S1), we compared average population synchrony across several conditions: (1) the bottom and top 20% of NREM trials ranked by frontal EEG slow-wave activity (0.5–4 Hz power) during baseline conditions ([-2,0]s relative to stimulus onset); (2) NREM trials from the first versus the last two hours of the recovery sleep period; and (3) different sleep stages and sleep deprivation states, based on a re-analysis of data from a previous study (Marmelshtein et al., 2023).

### Analysis of EEG slow-wave activity and auditory evoked potentials (Figures 6A-B, 7A-B)

Frontal EEG signals were extracted from a 2-second window surrounding click-train stimuli ([-1, 1] s). The auditory evoked potential (AEP) for 2- and 10-Hz click trains was obtained by averaging 100 consecutive trials relative to the proximate wake-sleep transition (Figures 6A-B, 7A-B). To measure the magnitude of the AEP’s negative deflection, we identified the minimum point between 50–150 ms after stimulus onset and calculated the integral of the area within this trough. Baseline (pre-stimulus) slow-wave activity (SWA; Figures 6B, 7B, diamonds) was estimated using the average Welch’s power spectral density of the EEG signal within the 0.5–4 Hz frequency range.

### Feature modulation between states (Figures 4D, 5D)

For each session, all auditory stimulation trials occurring around falling asleep (or awakening) episodes were sorted according to their time relative to the proximate moment of falling asleep (or awakening). We defined three vigilance intervals in the falling asleep process: “wakefulness” / “First seconds of NREM” / “NREM”, with different time ranges relative to the moment of falling asleep (-20s < t < -5s, 5s < t < 10s, 60s < t; respectively). Two intervals were also defined for awakening episodes: “Sleep” / “First seconds of wakefulness” (-20s < t < -5s, 5s < t < 10s; respectively). Trials around sleep onset / awakenings were sorted into these intervals.

For each feature of auditory processing (spontaneous firing, onset response, population synchrony, post-onset firing, 40 Hz locking), we pooled trials from all relevant click-trains presented (2,10,20,30,40 Hz): for spontaneous firing rate and population synchrony we examined baseline activity of all click trains; for onset firing rate, responses to all click trains; for post-onset firing rate click trains of 2 clicks/s and 10 clicks/s (allowing sufficient inter click interval to not interfere with the post-onset period); and for 40 Hz locking click trains of 40 clicks/sec.

For each neuronal response feature, the average value across all trials within each vigilance interval was calculated. To quantify modulation across wake-sleep transitions, we computed a modulation index between intervals as follows:

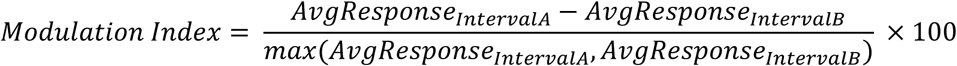

For the falling asleep process (Figure 4), we compared “first seconds of NREM” vs. “wakefulness” (Figure 4D, dark turquoise bars) and “NREM” vs. “wakefulness” (Figure 4D, black bars). For awakenings (Figure 5), we compared “first seconds of wakefulness” vs. “NREM” (Figure 5D, dark turquoise bars).

### Feature analysis along arousal state transitions (Figures 4C, 5C, 6B, 7B)

For each session and per each feature, auditory trials within falling asleep (awakening) episodes were extracted and sorted by time relative to the moment of falling asleep (awakening) as before, yielding a sorted unit response raster (Figure 4A). For each unit, the responses for adjacent trials (in terms of time relative to the proximate sleep onset/awakening instance) were smoothed by a moving average window (uniformly weighted), with the size of 50 trials and an overlap of 49 trials (figures 4-5). For EEG signals, a 100-trials window size was used with a 99-trials overlap (figures 6-7). Due to unevenly spaced stimuli’s relative times, new time stamps were created by smoothing the original stimuli relative times with the same method. Resulting in a modified raster where each row represents a 50-trial averaged response and its average stimuli time. Thereafter, a modulation index was calculated for each timepoint between each 50-trial (or 100-trial) average neuronal (EEG) response feature to that during the pre-transition interval (“wakefulness” for falling asleep, Figures 4C, 6B; and “Sleep” for awakening, Figures 5C, 7B). For population synchrony and EEG SWA, the feature values were first calculated for each trial, and then averaged using a running 50-trial window.

The continuous modulation index for each feature was averaged among all units within a session to gain the sessions’ mean continuous modulation index. Next, for each session and feature, the continuous index was interpolated, using the ‘interp1’ function with ‘pchip’ as the interpolation method (MATLAB, MathWorks), to gain evenly spaced and mutual values for all sessions. Thereafter, the continuous values were averaged into a grand mean of all sessions of continuous modulation index along the falling asleep (awakening) process (figures 4C, 5C, 6B, 7B).

### Temporal dynamics upon falling asleep analysis (Figures 4E-F, 6C-D)

For each session, trials were ordered based on their timing relative to the nearest sleep onset instance (as in Figure 4A). We extracted the following measures for each trial: post-onset firing rate (FR), population synchrony, spontaneous FR, and 40-Hz locking (n=7 sessions for post-onset firing, n=15 sessions for other features). To normalize per-unit responses, post-onset FR, spontaneous FR, and 40-Hz locking were divided by each unit’s mean response across all trials. Population synchrony, which represents a correlation coefficient naturally constrained between [-1,1], did not require normalization.

For each neuronal response feature, we computed a session-level population mean per trial by averaging values across all recorded units. To smooth fluctuations across trials, a moving average window of 50 trials (with an overlap of 49 trials) was applied. Since stimulus timings were unevenly spaced relative to sleep onset, new time stamps were generated using the same smoothing method. Both the neuronal response and time vectors were then fitted to a four-parameter sigmoid function using **LSQCURVEFIT.m** (MATLAB, MathWorks):

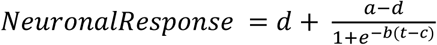

where *t* represents the new time stamps (mean trial time relative to sleep onset), *b* is the sigmoid slope steepness, *c* is the inflection point (midpoint), and *a* and *d* are the upper and lower asymptotes, respectively. Sessions with a good fit (R^2^>0.4) were included in the analysis and quantification in Figures 4E-F (n = 7, 10, 11, and 12 sessions for post-onset FR, population synchrony, spontaneous FR, and 40-Hz locking, respectively).

The sigmoid model parameters were used to define key transition points (Figure 4E, left): **Midpoint:** The sigmoid’s inflection point (c). **Onset and Offset:** Time points at 10% and 90% along the sigmoidal slope, calculated as:

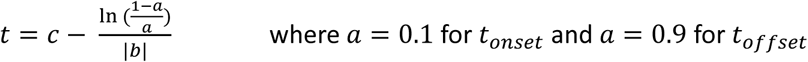

where *a* = 0.1 for *t*_*onset*_ and *a* = 0.9 for *t*_*offset*_ and **Duration:** Defined as *t*_*offset*_ − *t*_*onset*_.

To generate the normalized modulation traces in Figure 4E, we standardized sigmoid fits by setting *a* = 1, *d* = 0, and *b* = |*b*|, ensuring all curves ranged from [0,1] with a positive slope. This allowed direct comparisons of increases and decreases across measures, such as post-onset FR and population synchrony.

For EEG analysis (Figure 6C-D), we included only sessions with 2- and 10-Hz click trains (n = 7) relevant for calculating click-evoked auditory evoked potentials (AEPs). Frontal EEG baseline slow-wave activity (0.5-4 Hz power, measured from -1 to 0 s relative to click onset) was extracted per trial and smoothed using an 80-trial moving average window (overlap = 79 trials). EEG AEP negativity was quantified by first averaging click-evoked EEG signals over 80-trial groups (to obtain the AEP), identifying the minimum within the [50, 150] ms trough, and computing the integral of the trough area. Sigmoid fitting and calculation of onset, midpoint, offset, and duration for EEG measures followed the same procedure as neuronal data. All EEG sessions showed a sufficiently good fit (R^2^>0.4, n=7).

### Silent Intervals Analysis (Figure S1)

To identify potentially local silent intervals, we conducted our analysis on a per-microwire basis, aggregating spikes from all recorded clusters within each microwire. Silent intervals were defined as periods of neuronal silence lasting at least 50 ms (Vyazovskiy et al., 2009), and their probability was assessed in the post-onset window ([30,80] ms; Figure S1A). To account for changes in silent interval probability driven solely by fluctuations in spontaneous firing rates, we computed the absolute silent interval probability and subtracted the expected probability derived from a simulated Poisson-process unit with the same firing rate (Δ50 ms silence probability in Figure S1B).

### Rejecting non-significant units

Units with an average spontaneous firing rate <0.5 spikes/sec in any of the vigilance states were excluded from the analysis. Units without a significant response to a single-click stimulus were also excluded from our analysis. The significance of the response was calculated using the Monte Carlo Permutation Test, which was conducted as follows. A unit’s PSTH, in the time interval of [-500ms, 30ms] relative to the beginning of each 2 Hz and 10 Hz Clicks/sec trial, underwent minimal temporal smoothing (gaussian kernel of σ = 2). The significance was examined separately for the trials before the moment of falling asleep (or awakening) and for the trials after the moment of falling asleep (or awakening). Among the trials of each group, the PSTHs in the baseline interval ([-500ms,0] before click onset) were averaged, as well as the activity in the onset interval ([0, 30ms] after click onset). The difference between the maximum of the averaged PSTH in the time window of the onset and the baseline was calculated. After that, we shuffled between the baselines and the onsets of the smoothed PSTHs (that is, randomly, for 50% of the trials, an exchange was made between the time window of the baseline and the time window of the onset), and then again, we averaged over all the trials, and calculated the difference in the firing rate in the time windows of onset to baseline. This process was performed 1000 times to calculate the surrogate distribution from which the p-value of the real onset response magnitude was derived. Units whose response was not significant, for both before and after falling asleep (or awakening), were also excluded from the analysis.

### EEG analysis along arousal state transitions (Figures 6E-F, 7C-D, S3)

We utilized trial-by-trial variability to explore the relationship between the EEG and neuronal measures (Figures 6E-F, 7C-D, S3). As before, we temporally sorted all trials relative to the proximate falling asleep (or awakening) episodes. We then took groups of 16 consecutive trials and sorted them to four different groups - Low, Medium Low, Medium High, and High (L, ML, MH, H) - based on their relative post-onset response in the auditory cortex population (i.e. exactly four trials in each group). We then repeated this process for the next 16 trials, and so on, until we sorted all trials into one of these four groups. This process allowed us to create four groups of trials with gradually increasing levels of post-onset response (Figure 6Ei left), but with no statistically significant difference in their average timing relative to the proximate falling asleep (or awakening) episode (p=0.3916, n=7 sessions, Friedman test). We then calculated for each of the four groups of trials the average population synchrony (Figure 6Ei right), EEG AEP (Figure 6Eii left), AEP negative deflection magnitude (Figure 6Eii right), EEG power spectrum (Figure 6Eiii left) and EEG SWA (Figure. 6Eiii right).

In Figure 6F we performed the same analysis as in Figure 6E described above, but instead of sorting trials based on their relative post-onset response, we instead sorted them based on the relative population synchrony. This (by definition) created four groups of trials with increasing levels of population synchrony (Figure 6G right), but with no statistically significant difference in their average timing relative to the proximate falling asleep (or awakening) episode (p=0.7736, n=7 sessions, Friedman test). In Figure S3A and S3B we applied the same analysis but with sorting trials based on their relative onset FR and Spontaneous FR, respectively.

To analyze spectral power changes in the EEG induced by click trains (Figure S2), we applied the event-related time-frequency decomposition function newtimef.m (EEGLAB (Delorme and Makeig, 2004)) to frontal EEG responses to 2- and 10-Hz click trains. The analysis used a fixed 500 ms temporal window, a baseline period of [-1,0] s relative to stimulus onset, and averaged power changes within the [500,1500]s temporal window for each frequency (Figure S2B).

### Statistics

We used a linear mixed-effects model (LME) to analyze electrophysiological neural data due to its nested and hierarchical nature (Aarts et al., 2014; Makin and De Xivry, 2019). This model was necessary to account for non-independencies in measures from different units obtained in the same electrode, experimental session, or animal. We included animal identity as a random effect, as well as experimental session and microwire electrode as nested random effects within each animal. We obtained model parameters using the ‘fitlme’ function (Matlab, MathWorks) and utilized restricted maximum likelihood estimation.

When presenting average effects per session, we only considered sessions that had a minimum of 3 units. Throughout the main text, all effect sizes are presented as mean±SEM for all units unless specified otherwise. To evaluate variance across different conditions (>2), we conducted a Friedman test (similar to a non-parametric repeated measures ANOVA) on data summarized at the animal level. This involved averaging all the units for each of the 6 animals. To evaluate differences in EEG activity across two conditions (e.g., comparing wakefulness vs. first-seconds NREM) we conducted a Wilcoxon sign-rank test (WSRT). To evaluate the temporal dynamics for different neuronal and EEG measures (Figures 4E-F, 6C-D, onset, midpoint, offset and duration of sigmoid fit), each with a different number of sessions, a Wilcoxon Rank-Sum test (WRST) was conducted. For the EEG measures (AEP negativity and SWA) obtained from the same 7 sessions, a paired WSRT showed similarly significant results (p=0.0156 for midpoint, offset and duration, n=7).

## Acknowledgements

We thank Dr. Noa Bar-Ilan Regev for administrative assistance. Supported by the Israel Science Foundation (ISF) grants 1326/15 and 1557/22 and by the European Research Council grant ERC-2019-CoG 864353. A.M. is supported by the EMBO ALTF 486-2024 and Rothschild fellowship.

## Author contributions

A.M. and Y.N. designed the project. A.M. built experimental setup, compiled stimuli, and performed experiments. B.L. led data analysis with support from A.M. and B.H. Y.N. supervised the project and secured funding. All authors wrote the paper.

