## Supplementary Material for "Abrupt and gradual changes in neuronal processing upon falling asleep and awakening"

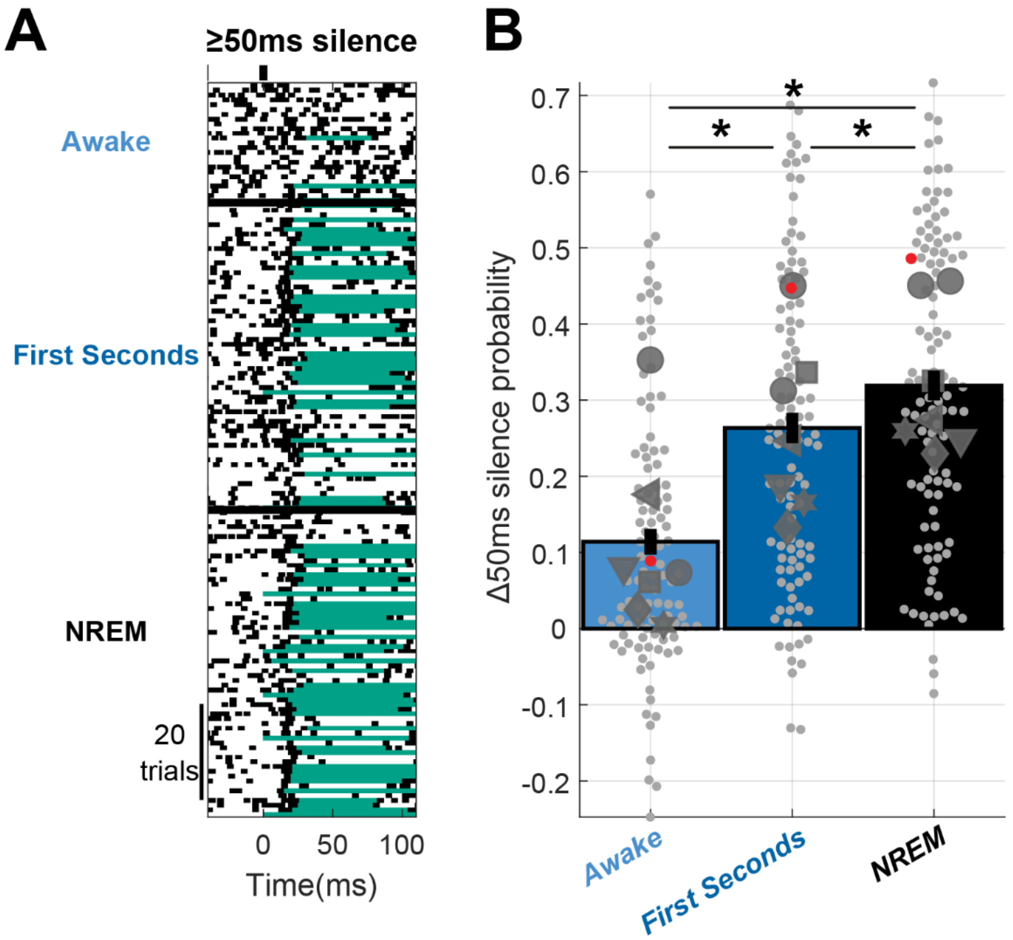


**Figure S1. Stimulus-induced silent intervals emerge within first seconds of falling asleep.** (A) Representative multi-unit raster in response to a click while the animal was awake, in the first seconds of NREM sleep, and NREM sleep >60 seconds after the moment of falling asleep. Silent intervals (≥50ms of no firing) spanning the entire post-onset period ([30, 80]ms) are marked in green. (B) Increase in silent intervals probability (relative to Poisson process) across awake, first seconds and NREM sleep conditions, for all electrodes (n = 105) and sessions (n = 7). Bars and error bars represent mean±sem of all electrodes. Small gray markers represent individual electrodes (red for representative electrode in A). Large dark gray markers represent individual session mean (with each shape representing an individual animal). *$p < 0.05$. mean±sem increase in probability 0.11±0.042, 0.26±0.039 and 0.32±0.033 for “wakefulness”, “first seconds of NREM” and “NREM”, respectively.

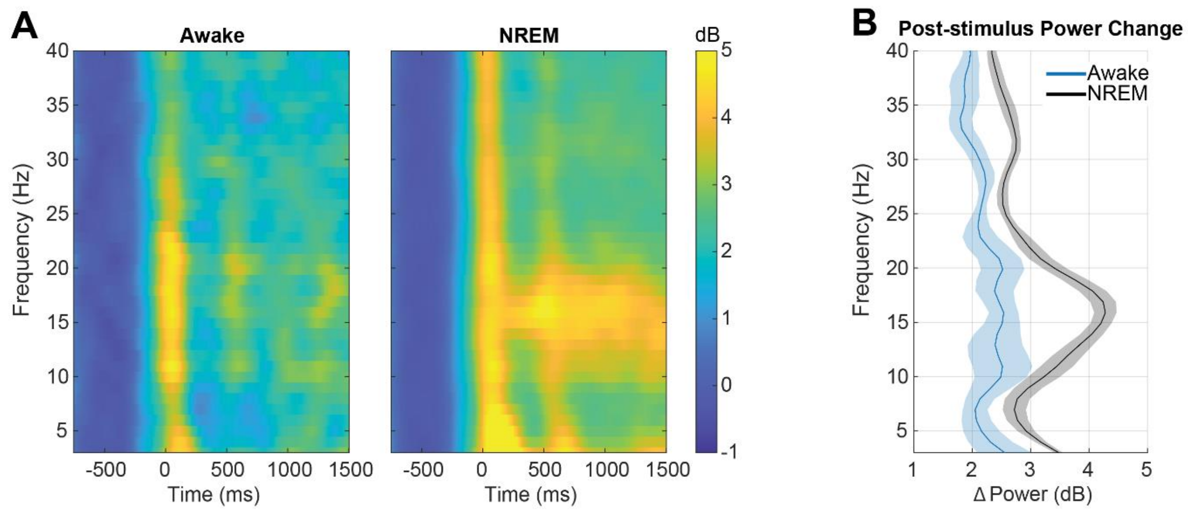


**Figure S2. Enhanced EEG spindle frequency power following a click stimulus during NREM sleep.** (A) Spectrogram representing average (n=7 sessions) induced EEG power changes following a click (0ms time) before and after falling asleep ("awake": at least 5s before falling asleep; "NREM": at least 5s after falling asleep). (B) Average power change from baseline at the [500,1500]ms peri-stimulus interval during wakefulness and NREM sleep (n=7 sessions). Traces and shadings represent mean±sem. A power increase in the spindle frequency range occurs only during NREM sleep.

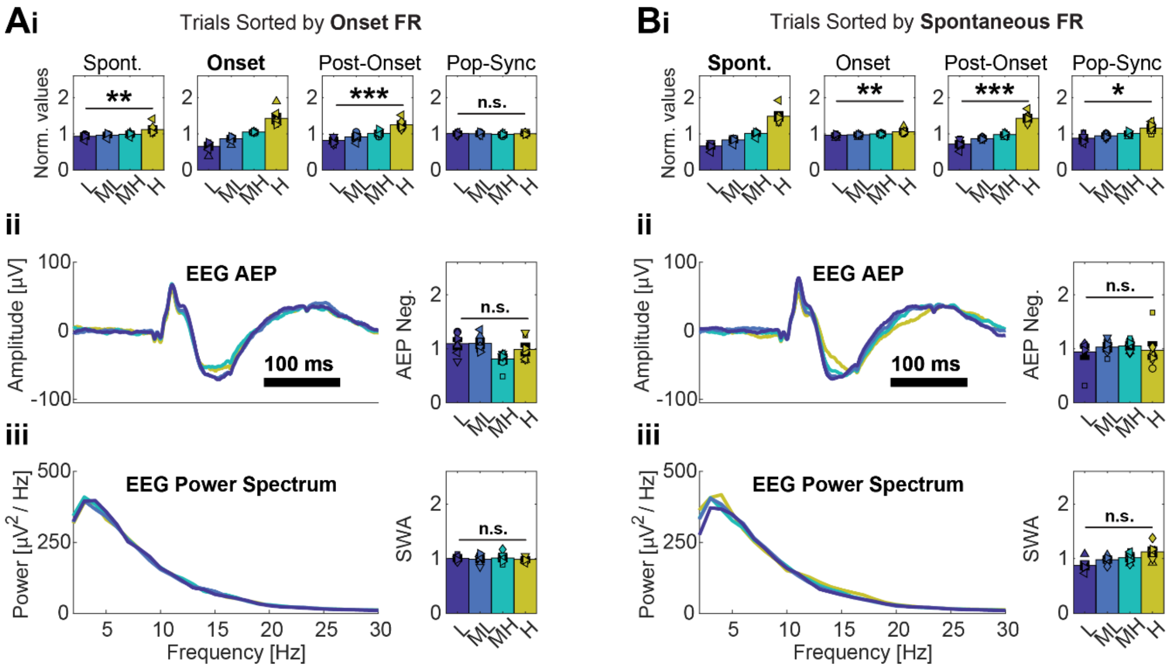


**Figure S3. Onset and spontaneous firing while falling asleep are not associated with EEG AEP or SWA.** (A,B) Same as Figure 6E,F but with trials sorted according to onset and spontaneous FR. (A) trials were sorted into four quartiles based on their relative auditory cortex onset firing (low onset FR (L), medium-low (ML), medium-high (MH) and high(H). (Ai) Auditory cortex normalized spontaneous, onset and post-onset firing and population synchrony (left to right) averaged across all sessions (n=7) for the four different trial groups. (Aii) Average frontal EEG AEP (left) and its negative deflection magnitude (right) for the four different trials groups across all sessions. (Aiii) Average baseline (pre-stimulus) power spectrum (left) and slow-wave activity [SWA] (right, 0.5-4 Hz power) for the four different trials groups across all sessions. (B) same as A but with trials sorted into four quartiles based on their relative spontaneous firing in auditory cortex (low spontaneous FR (L), medium-low (ML), medium-high (MH) and high(H)). Both neuronal measures (spontaneous and onset firing) were not associated with either of the EEG measures (AEP negative deflection and baseline SWA).

| State | Population Synchrony | Dataset |
|---|---|---|
| vigilant wakefulness | 0.17±0.006 | Modulation by sleep deprivation and sleep stages. N=324 units. reanalysis of data from (Marmelshtein et al., 2023) |
| wakefulness after 5h sleep deprivation | 0.21±0.007 | |
| recovery REM sleep | 0.2±0.007 | |
| recovery NREM sleep | 0.33±0.009 | |
| bottom 20% SWA trials | 0.2±0.009 | Modulation by EEG slow-wave activity during NREM sleep. N=151 units. |
| top 20% SWA trials | 0.25±0.012 | |
| first 2h of recovery sleep period | 0.24±0.01 | Modulation by sleep pressure during NREM sleep. N=151 units. |
| last 2h of recovery sleep period | 0.21±0.01 | |

**Table S1. Population synchrony depends on sleep deprivation, sleep stages, EEG slow-wave activity and sleep pressure.** Analysis of data from a previous publication (Marmelshtein et al., 2023) and current datasets demonstrating that population synchrony captures the synchrony characteristics of sleep. Values represent the mean±SEM of the Pearson correlation coefficient of each unit's firing rate (in 50ms bins) to that of the entire auditory cortex population ("population coupling" (Okun et al., 2015)).

| | Spontaneous firing (n=11 sessions) | Post-onset firing (n=7 sessions) | Population synchrony (n=10 sessions) | 40Hz locking (n=12 sessions) |
|---|---|---|---|---|
| Onset | -8.61±3.01s | -5.23±1.68s | -4.50±3.41s | -9.74±1.77s |
| Midpoint | -3.69±1.46s | -1.27±1.04s | 7.01±1.59s | 0.03±0.97s |
| Offset | 1.23±2.96s | 2.69±1.23s | 18.51±2.35s | 9.80±3.37s |
| Duration | 9.84±5.21s | 7.92±2.08s | 23.02±4.91s | 19.54±5.02s |

**Table S2**. **Comparison of the temporal dynamics of the four neuronal measures showing changes upon falling asleep.** Values represent the mean ± SEM timing of the “onset”, “midpoint” and “offset” of the sigmoid fit (10%, 50% and 90% along the sigmoid slope), as well as the duration (time difference between the onset and offset) across sessions (n = 7, 10, 11, and 12 sessions for post-onset FR, population synchrony, spontaneous FR, and 40-Hz locking, respectively).
